# Complement receptor *C3ar1* deficiency does not alter brain structure or functional connectivity across early life development

**DOI:** 10.1101/2025.04.24.650541

**Authors:** Hanna Lemmik, Eugene Kim, Eilidh MacNicol, Davide Maselli, Michel Bernanos, Zhuoni Li, Dauda Abdullahi, Esther Walters, Maria Elisa Serrano Navacerrada, Wuding Zhou, Aleksandar Ivetic, Diana Cash, Laura Westacott

## Abstract

Genetic deletion of the complement C3a anaphylatoxin chemotactic receptor (*C3ar1*), a key component of the innate immune response, is reported to induce behavioural phenotypes consistent with psychiatric symptomatology in mice, but when and where *C3ar1* is needed in the brain is unresolved. These questions are significant because, as a G-protein-coupled receptor, human C3AR1 serves as a potential therapeutic target for disorders associated with complement dysregulation, such as schizophrenia. To provide a brain-wide (where) assessment of developmental *C3ar1* activity, we used longitudinal (when) tensor-based morphometry, diffusion-weighted magnetic resonance imaging (MRI) and blood oxygen-level dependent functional MRI in male and female *C3ar1*-deficient mice and wild-type littermates, with behavioural assessment in adulthood. Unexpectedly, we did not find a robust *C3ar1*-dependent phenotype in any of these measures. Therefore, our study does not support neurodevelopmental hypotheses for *C3ar1*, which is encouraging for therapeutic strategies targeting this receptor since interventions are unlikely to disrupt brain development.

## Introduction

The complement system is a conserved immune pathway that participates in host defence through pathogen clearance and regulating inflammation (Chaplin 2020) as well as tissue homeostasis (Kunz and Kemper 2021; West and Kemper 2023), with emerging roles in neurodevelopment, psychiatric disorders and neurodegeneration (Stevens et al. 2007; Hong et al. 2016; Sekar et al. 2016).

The most convincing evidence for the involvement of the complement system in neurodevelopment so far is its genetic association with schizophrenia. Schizophrenia is a complex and highly heritable neurodevelopmental disorder characterised by hallucinations, delusions, and impaired cognition, with symptoms typically emerging in late adolescence or early adulthood (McCutcheon, Reis Marques, and Howes 2020; Howes, Bukala, and Beck 2024). Neurobiological hallmarks of schizophrenia include grey matter loss (Vita et al. 2012) and a reduction in synaptic density (Osimo et al. 2019). Genome-wide association studies (GWAS) of schizophrenia have helped to identify two complement-related risk loci; a structural variant in the complement component 4 A (*C4A*) gene (Sekar et al. 2016), which encodes the C4A protein responsible for propagation of complement activation, and a variant in the CUB and Sushi Multiple Domains 1 (*CMSD1*) gene, which encodes a putative complement inhibitor protein (Schizophrenia Working Group of the Psychiatric Genomics Consortium 2014; Baum et al. 2024). Preclinical studies link these variants to increased brain-specific complement activation and synapse loss (Sekar et al. 2016; Yilmaz et al. 2021; Baum et al. 2024), potentially tying complement activation to synaptic pathology in schizophrenia. Indeed, the *C4A* schizophrenia-risk genotype associates with MRI markers of grey matter loss and reduced cognitive performance in humans, even in the absence of neurological disorders (O’Connell et al. 2021).

Further, patients with elevated complement proteins have more severe negative symptoms in psychosis (Byrne et al. 2024), which typically do not respond to anti-psychotic medication. Complement modulation therefore has potential in addressing this unmet therapeutic need.

C3a anaphylatoxin chemotactic receptor (C3aR1), a G-protein coupled receptor (GPCR) bound by complement activation product C3a and the granin family neuropeptide TLQP-21 (Rodriguez et al. 2023), acts downstream of complement activation and stands out as a pharmacologically tractable target for modifying complement activity in the brain (Hauser et al. 2017). In mice, *C3ar1* transcripts are predominantly expressed by microglia, with minimal neuronal expression observed in both healthy adult humans and mice according to single-cell transcriptomic analyses (Hammond et al. 2019; Tasic et al. 2016). While its temporal expression patterns are not yet well characterised, C3aR1 appears to be active during early embryonic development, potentially influencing progenitor cell proliferation (Coulthard et al. 2018; Bénard et al. 2008; Hammond et al. 2019). C3aR1 also appears to facilitate developmental astrocyte phagocytosis by microglia in the retina (Gnanaguru et al. 2023), as well as to regulate microglial reactivity and neuroinflammation more broadly (Gedam et al. 2023; Ge, Guan, and Wang 2024; Zheng et al. 2021; Lian et al. 2015; Vasek et al. 2016; Chew and Petretto 2019). Brain morphological changes observed in *C3ar1*-deficient mice further support its neurodevelopmental relevance (Westacott et al. 2021; Pozo-Rodrigálvarez et al. 2021), although there is no consensus on the precise neurodevelopmental actions of C3aR1. Addressing this gap is important given this receptor’s potential as a pharmacological target.

C3aR1 signalling may impact brain functions relevant to psychiatric symptomatology since a range of behavioural phenotypes have been reported in *C3ar1*-deficienct mice. These include abnormal anxiety-like behaviours (Westacott et al. 2022), hyperactivity (Pozo-Rodrigálvarez et al. 2021), cognitive impairment (Coulthard et al. 2018) but also a resilience to depressive-like behaviours induced by chronic stress or inflammation (Crider et al. 2018; Zhang et al. 2022; Sun et al. 2024). Although the observed involvement of C3aR1 in behaviour suggests that it is needed for normal brain function, previous studies have not addressed the question of whether these phenotypes arise because of a C3aR1 deficit during development or because it is continuously needed, which can be resolved through longitudinal assessment. Another important yet overlooked aspect is the use of the appropriate wild-type *littermate* control animals that none of the aforementioned studies included. Instead, these studies used separately raised cohorts of control and mutant mice which represents a known confound due to the rapidly diverging genetic backgrounds in small inbred colonies (Fitch and Atchley 1985), but also because litter environment affects behaviour and brain development (Crews et al. 2009; Jiménez and Zylka 2021; Valiquette et al. 2023).

To investigate the potential consequences of *C3ar1* deficiency during development, we adopted a global, unbiased approach, conducting a longitudinal study of male and female *C3ar1*-deficient mice and their wild-type littermates during adolescence (postnatal day, PND30) and adulthood (PND90). Using structural and diffusion magnetic resonance imaging (MRI and dMRI), as well as resting state fMRI (rsfMRI), we aimed to assess whether the absence of C3aR1 affects brain development during adolescence—a critical period for psychiatric vulnerability (Westacott and Wilkinson 2022; Paus, Keshavan, and Giedd 2008)—or whether this requirement only becomes evident by adulthood.

Our imaging measures included regional brain volumes derived from tensor-based morphometry (TBM), fractional anisotropy (FA) from dMRI to evaluate white matter organisation, which is influenced by microglial activity during development (Chan et al. 2024; Falangola et al. 2023), and graph theoretical analysis of functional connectivity (FC) correlates of blood oxygenation level dependent (BOLD) signal, such as global efficiency and clustering coefficient to characterise brain network topology (Zhu et al. 2017; Forlim et al. 2024; Hadley et al. 2016). These techniques were complemented by behavioural testing in adult mice that measured cognition and emotional reactivity, as we sought to replicate previously reported behavioural experiments in *C3ar1*-deficient mice (Pozo-Rodrigálvarez et al. 2021; Westacott et al. 2022; Coulthard et al. 2018). Unexpectedly, we found no robust brain or behavioural phenotype in our datasets, challenging the previous assumptions of a neurodevelopmental role for C3aR1 under physiological conditions.

## Results

### *C3ar1*^tm1Cge^ homozygous mice lack detectable *C3ar1* mRNA

We used an established *C3ar1* mutant line, *C3ar1^tm1Cge^* (Humbles et al. 2000) and performed our own validation of the mutation. For this purpose, we designed a 79-base pair (bp) amplicon targeting the putatively deleted region (**Figure 1a**). PCR analysis of cDNA derived from bone marrow-derived macrophages—a cell type consistently reported to express high levels of *C3ar1* mRNA (Tao et al. 2021; Mommert et al. 2018; Mamane et al. 2009)—showed no detectable transcript in this region in the mutant animals (**Figure 1c**), confirming the absence of the canonical transcript. The *C3ar1* gene contains an in-frame start codon after the deletion (**Supplemental figure 1)**. We also confirmed that no transcript is made that includes this region (**Figure 1b**). These results were further corroborated by quantitative PCR (qPCR, n = 8) in homozygous knockout animals in either M0-like or interleukin 4 (IL4)-induced M2-like macrophages (**Supplemental figures 2a–e**). Together, these results confirm that *C3ar1^tm1Cge^* is a true loss-of-function or a “knockout” allele resulting in no detectable *C3ar1* transcript.

**Figure 1.**
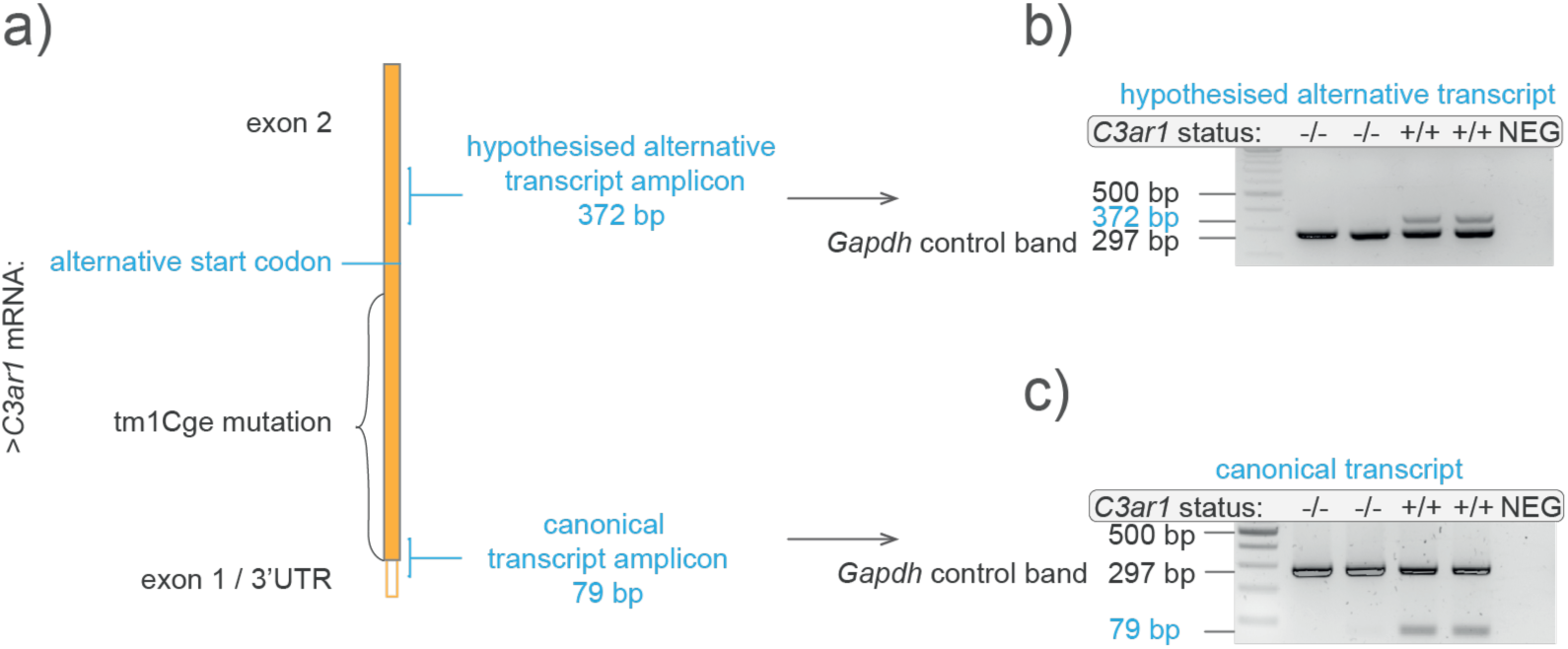
*C3ar1*^tm1Cge^ homozygous mice lack detectable *C3ar1* mRNA. **(a)** Schematic shows *C3ar1* mRNA with its only protein coding exon and the design of the 79 bp PCR amplicon which targets an exon-1/5’ untranslated region (UTR)-exon-2 junction of the canonical *C3ar1^+/+^* transcript. Also shown is another 372 bp amplicon which targets a region downstream of the deletion and after an alternative start codon. The start of exon 2 is deleted in *C3ar1^-/-^* mice, so PCR should result in no amplification. Similarly, if no alternative transcript is present, there should be no amplification**. (b–c)** Gel images show PCR products of cDNA from bone-marrow derived interleukin 4 (IL4)-induced M2-like macrophages. Each sample well has a 297 bp Glyceraldehyde 3-phosphate dehydrogenase (*Gapdh*) control band. NEG: no reverse transcriptase negative control carried over from cDNA synthesis. **(a)** Canonical transcript. **(b)** Hypothesised alternative transcript. The bands appear hollow due to over-saturation. -/- = *C3ar1^-/-^*, +/+ = *C3ar1^+/+^*.

### *C3ar1*-deficiency does not influence total or regional brain volume

We conducted a longitudinal MRI study (referred to as Cohort 1 hereafter or implied when cohort is not specified) to investigate potential genotype-related differences in brain structure and function using *C3ar1^tmCge^* homozygous knockout mice (*C3ar1^-/-^*, *C3ar1*-deficient) and their littermate wild-type controls (*C3ar1^+/+^*) (Figure 2a) on C57BL6J (Charles River, UK) background. Both groups underwent *in vivo* MRI in adolescence (PND27–31) and adulthood (PND81–92). Structural MR images were additionally collected *ex vivo* from the same mice sacrificed in adulthood immediately after the *in vivo* scan to achieve higher isotropic resolution (0.1 mm *ex vivo* vs 0.15 mm *in vivo*) and increased signal-to-noise ratio (SNR). To corroborate these results, we also conducted MRI in adulthood (PND74-110) in an independent study cohort, referred to as Cohort 2.

**Figure 2.**
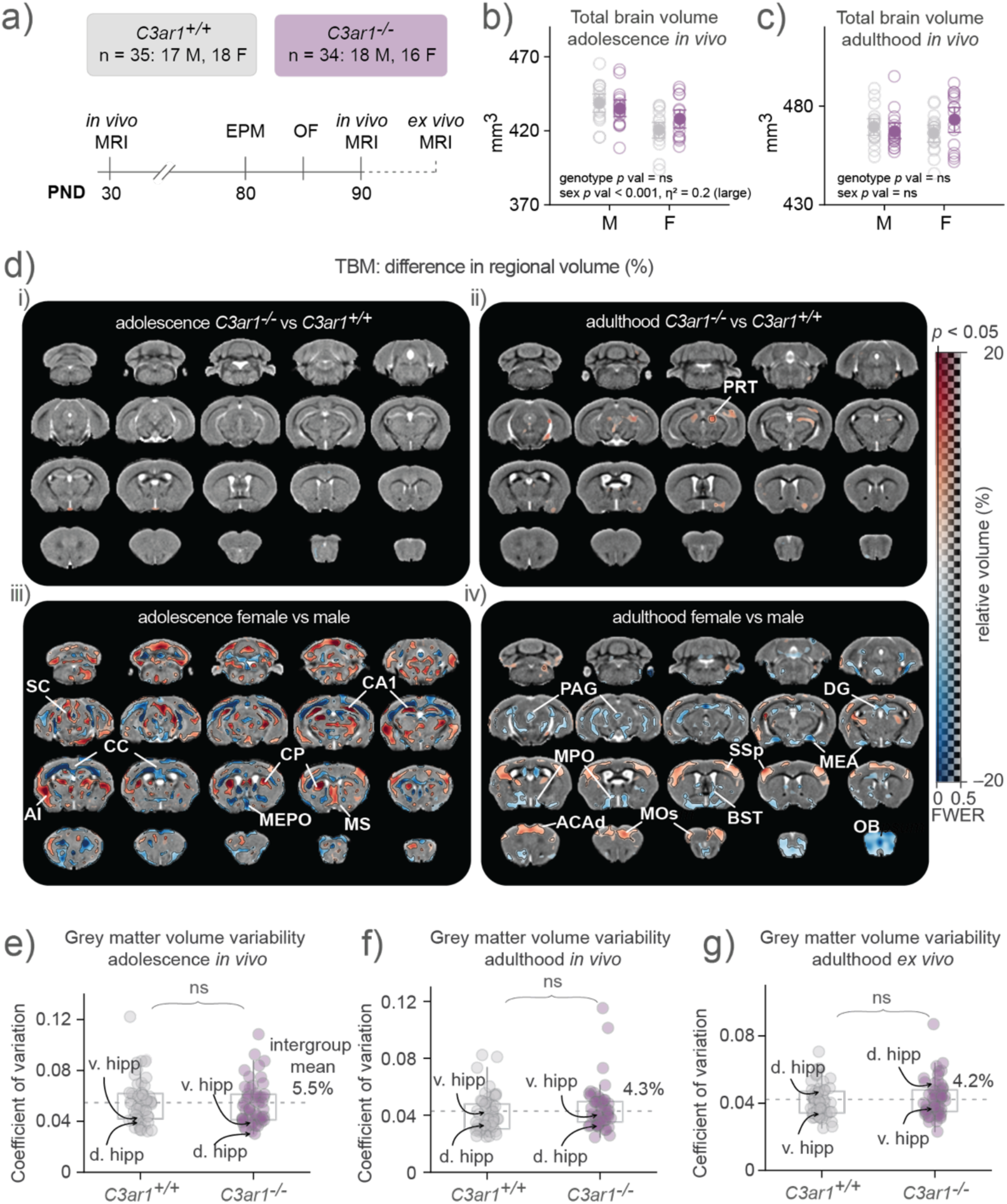
Sex but not *C3ar1* status influences regional brain volume. **(a)** Schematic of the MRI study shown in **(d–g)**. 69 mice were scanned twice *in vivo*; in adolescence at ∼ postnatal day (PND) 30 (range 27-31) and at adulthood ∼ PND90 (range 81-92), as well as once *ex vivo* after sacrifice after the adulthood scanning session. Behavioural tests were carried out before the adulthood scanning session. EPM = elevated plus maze, OF = open field. **(b)** Total brain volume in adolescence. Two-way ANOVA, genotype, sex, genotype * sex, F_[1,65]_ = 0.35, 16.72, 3.26, *p* = 0.56, <0.001, 0.76. **(c)** Total brain volume in adulthood. Two-way ANOVA, genotype, sex, genotype * sex, F_[1,65]_ = 0.74, 0.26, 3.33, *p* = 0.39, 0.61, 0.07. **(b–c)** Data presented as mean ± 95% CI. **(d)** Panels showing relative regional volume changes (%) overlaid on study-specific coronal templates (grey). Red hues signify areas larger in *C3ar1****^-/-^* (i–ii)** or females **(iii–iv)** and blue hues signify areas larger in *C3ar1^+/+^* **(i–ii)** or males **(iii–iv)**. Transparency of the colour overlay shows the statistical significance, ranging from family wise error (FWE)-corrected *p* value 0.5 to 0 (transparent to opaque, respectively). Areas where FWE-corrected *p* value < 0.05 are demarcated with a black line, and where the *p* value > 0.5 are grey (no overlay), meaning that in adolescence genotype comparison **(i)**, no voxels had a *p* value < 0.5. The locations of the coronal slices in relation to bregma in the left most column from top: −7.6, −4.6, −1.6, 1.4 mm. *C3ar1^+/+^* n = 35, 17 males and 18 females; *C3ar1^-/^*^-^ n = 34, 18 males and 16 females. ACAd = dorsal anterior cingulate cortex, AI = agranular insular cortex, BST = bed nucleus of stria terminalis, CA = cornu ammonis, CP = caudoputamen, DG = dentate gyrus, MEA = medial amygdala, MEPO = medial preoptic nucleus, MOs = secondary motor cortex (MOs), MPO = medial preoptic area, MS = medial septum, OB = olfactory bulb, PAG = periaqueductal grey, PRT = pretectal area, SC = superior colliculus, SSp = primary somatosensory cortex. **(e)** Coefficient of variation (CV) in adolescence *in vivo* for 72 regional volumes across all animals in the experiment. **(f)** CV in adulthood *in vivo*. **(g)** CV in adulthood *ex vivo*. **(e–g)** Data are shown with quartiles and whiskers show the extent of the distribution. Dotted line shows the intergroup mean excluding cerebrospinal fluid (CSF) areas. Kruskal Wallis *p* values, all ns. D. hipp = dorsal hippocampus, v. hipp = ventral hippocampus.

We used tensor-based morphometry (TBM) analysis to estimate total brain volume and to map regional brain volume differences. There were no genotype-dependent differences in total brain volume *in vivo* in adolescence (Figure 2b) nor in adulthood (Figure 2c). In all adolescent mice, irrespective of genotype, female mice exhibited significantly smaller total brain volumes compared to males. These sex differences disappeared by adulthood—a finding that aligns with previous observations in wildtype mice (Guma et al. 2024).

Still using TBM, no significant genotype-dependent differences in regional brain volumes were detected in adolescence (Figure 2d-i). In adulthood (Figure 2d**-ii**), *C3ar1^-/-^* mice showed a significant (*p* < 0.05) volume increase in the right pretectal area, and subthreshold (0.05 < *p* < 0.5) increases in the left pretectal area and the right lateral thalamus *in vivo*, but these differences were no longer observed in the same study cohort *ex vivo* despite improved spatial resolution (not shown since these group-level data would be presented as empty study-template maps), nor did we observe any genotype-dependent differences using TBM analysis in Cohort 2 (not shown). We also did not observe any significant genotype-by-sex interaction effects (not shown).

While no reproducibly significant genotype effects in regional volumes were observed in Cohort 1, we nevertheless detected sexually dimorphic effects (Figure 2d**-iii** and **2d-iv**). Adolescent female mice had significantly larger volumes (relative to total brain volume) bilaterally in the agranular insular cortex (AI), superior colliculus (SC), medial septum (MS) and in the CA1 region of the hippocampus, whereas male mice had increased relative volumes in white matter areas including the olfactory tract, corpus callosum, hippocampal commissure, and notably also in the median preoptic nucleus (MEPO) which is well known to be larger in male rodents (Gorski et al. 1978) (Figure 2d**-iii, Supplemental** figure 3 for absolute volume).

Echoing total brain volume measures, many of these sex differences were no longer observed in adulthood (see also **Supplemental** figure 4 for Cohort 2 data), but females showed larger relative volumes in the dorsal anterior cingulate cortex (ACAd), secondary motor cortex (MOs) and primary somatosensory cortex (SSp). Adult males had larger relative volumes in areas including the medial amygdala (MEA) and the bed nucleus of stria terminalis (BST) which, like the MEPO, are previously documented sexual dimorphisms (Hines, Allen, and Gorski 1992) and which were also observed *ex vivo* in this study cohort (**Supplemental** figure 5).

There were no genotype nor sex-by-genotype interactions on regional brain volumes at either time-point (not shown). Overall, female somatosensory and motor cortices increased more in volume between adolescence and adulthood than male (**Supplemental** figure 6), in line with the observed smaller differences in these areas in adulthood compared to adolescence.

If variability was high within our study sample, particularly in the *C3ar1*-deficient group, it would have hampered our ability to detect significant differences. The coefficient of variation for grey matter region-of-interest (ROI) volume showed no differences between genotypes *in vivo* in adolescence (Figure 2e) or adulthood (Figure 2f), nor *ex vivo* in adulthood (Figure 2g). The overall variability was low, ranging from 4.2–5.5% *in vivo* and 4.2% *ex vivo*, while hippocampal variability which has previously been repeatedly measured establishing a neuroimaging gold standard at 5% variability (Lerch et al. 2012), was 3.7–3.8% *in vivo* and 4.2% *ex vivo* in our study. These values suggest that our study was well-positioned to detect a genotype effect if one had been present.

### *C3ar1*-deficiency does not influence fractional anisotropy

To assess the potential impact of *C3ar1* deficiency on white matter integrity, we measured fractional anisotropy (FA) as an indirect marker of axonal microstructure (Friedrich et al. 2020) (Figure 3). Using voxel-wise analysis, we observed sub-threshold, non-significant (0.05 < *p* < 0.5) decreases in FA in *C3ar1*-deficient mice compared to wild-types (Figure 3a). In adolescence, sub-threshold reductions were noted in the corpus callosum (CC) and optic tract (OPT, Figure 3a, upper panel). In adulthood, they were primarily localised to the CC (Figure 3a, bottom panel). These sub-threshold differences were not observed *ex vivo*. Compared to males, female mice had sub-threshold decreases in FA *in vivo* in adulthood in the internal capsule and in the third ventricle (**Supplemental** figure 7), but this was again not observed *ex vivo* (not shown). We did not observe any sex-by-genotype interactions in our FA datasets (not shown).

**Figure 3.**
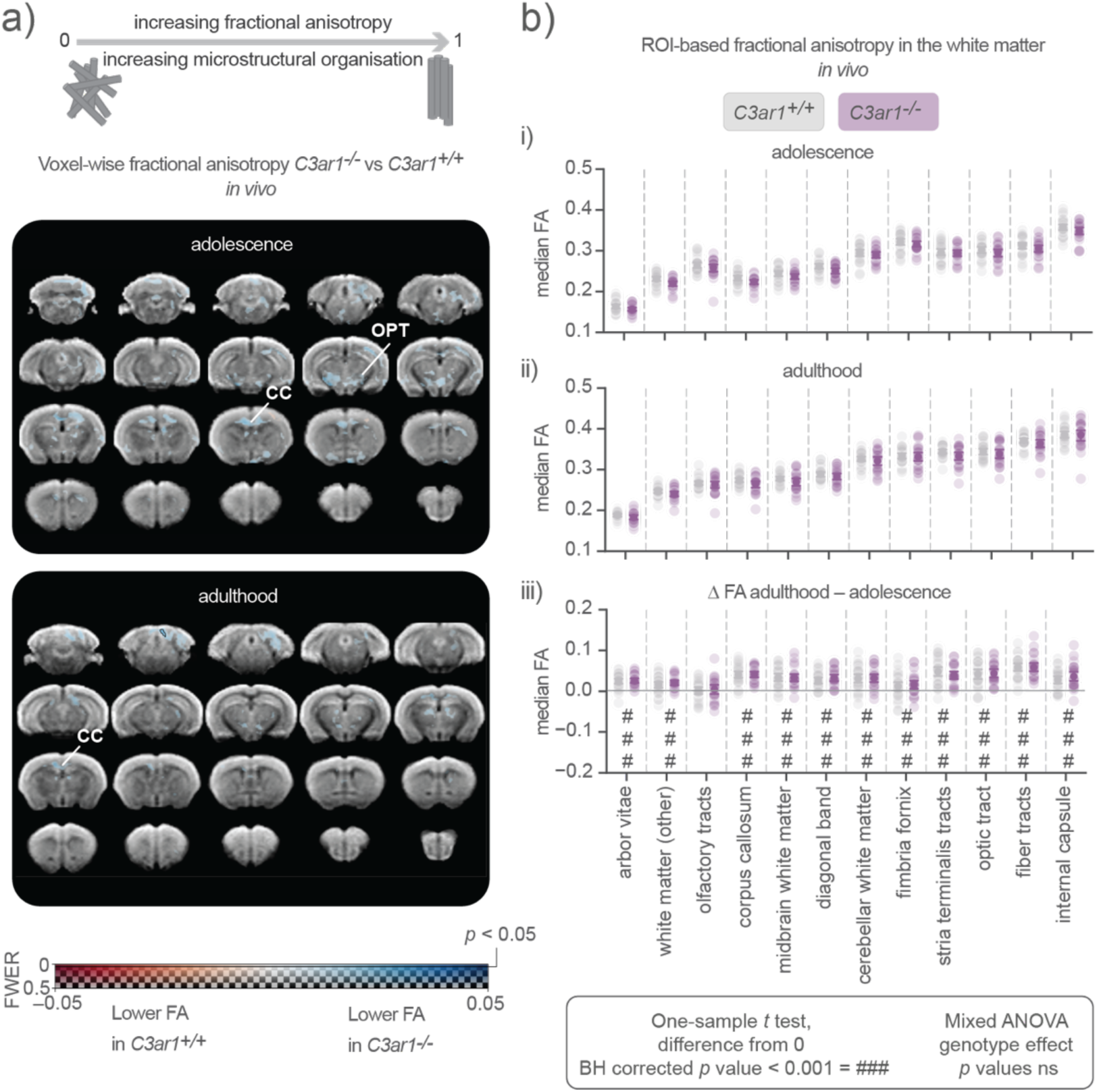
Fractional anisotropy does not depend on *C3ar1* status but increases with age. **(a)** Panels showing voxel-wise fractional anisotropy analysis in adolescence (top) and adulthood (bottom) corrected for FWE. There are no significant voxels (no black contour) where genotype effect is significant (*p* < 0.05). CC = corpus callosum, OPT = optic tract. **(b)** *In vivo* ROI-based median fractional anisotropy (FA) values in white matter regions for **(i)** adolescence, **(ii)** adulthood and **(iii)** change over time (adulthood – adolescence). Two-sided one-sample *t* test (difference from 0) *p* values that were adjusted with Benjamini-Hochberg (BH) procedure (### *p* value < 0.001). **(b-i–iii)** Mixed ANOVA with genotype and genotype-by-region interaction effects, all ns. Data are presented as mean ± 95% CI.

In our ROI-based FA analysis, we focused on predefined white matter regions, hypothesising that FA alterations would primarily occur in areas containing axonal tracts due to reported increased microglial phagocytosis in *C3ar1*-deficient mice during development (Gnanaguru et al. 2023; Falangola et al. 2023; Chan et al. 2024). In this analysis also, no significant genotype effects were observed in adolescence (Figure 3b-i) or adulthood (Figure 3b**-ii**). There were no genotype-dependent differences in the change in FA over time, but FA increased between adolescence and adulthood in all white matter areas except the olfactory tract (Figure 3b**-iii**), which is in line with the reported early maturation of the olfaction system in mice (Gretenkord et al. 2019).

### *C3ar1*-deficiency does not influence global functional brain connectivity

To evaluate global brain functional connectivity (FC), which has been reported to be altered in other mice who lack a specific microglial gene (Filipello et al. 2018; Deivasigamani et al. 2023; Zhan et al. 2014), we estimated FC through analysis of BOLD signal time-courses with the assumption that the magnitude of correlation between these time-courses corresponds to the strength of FC. We calculated pairwise correlation coefficients between 36 (18 per hemisphere) grey matter regions. Non-zero correlation values were averaged across proportional progressively decreasing sparsity thresholds to preserve biologically meaningful weak correlations while minimising noise (Bassett et al. 2009). No genotype-dependent differences in global FC were observed in adolescence (Figure 4a), adulthood (Figure 4d) or in the change over time (Figure 4g).

**Figure 4.**
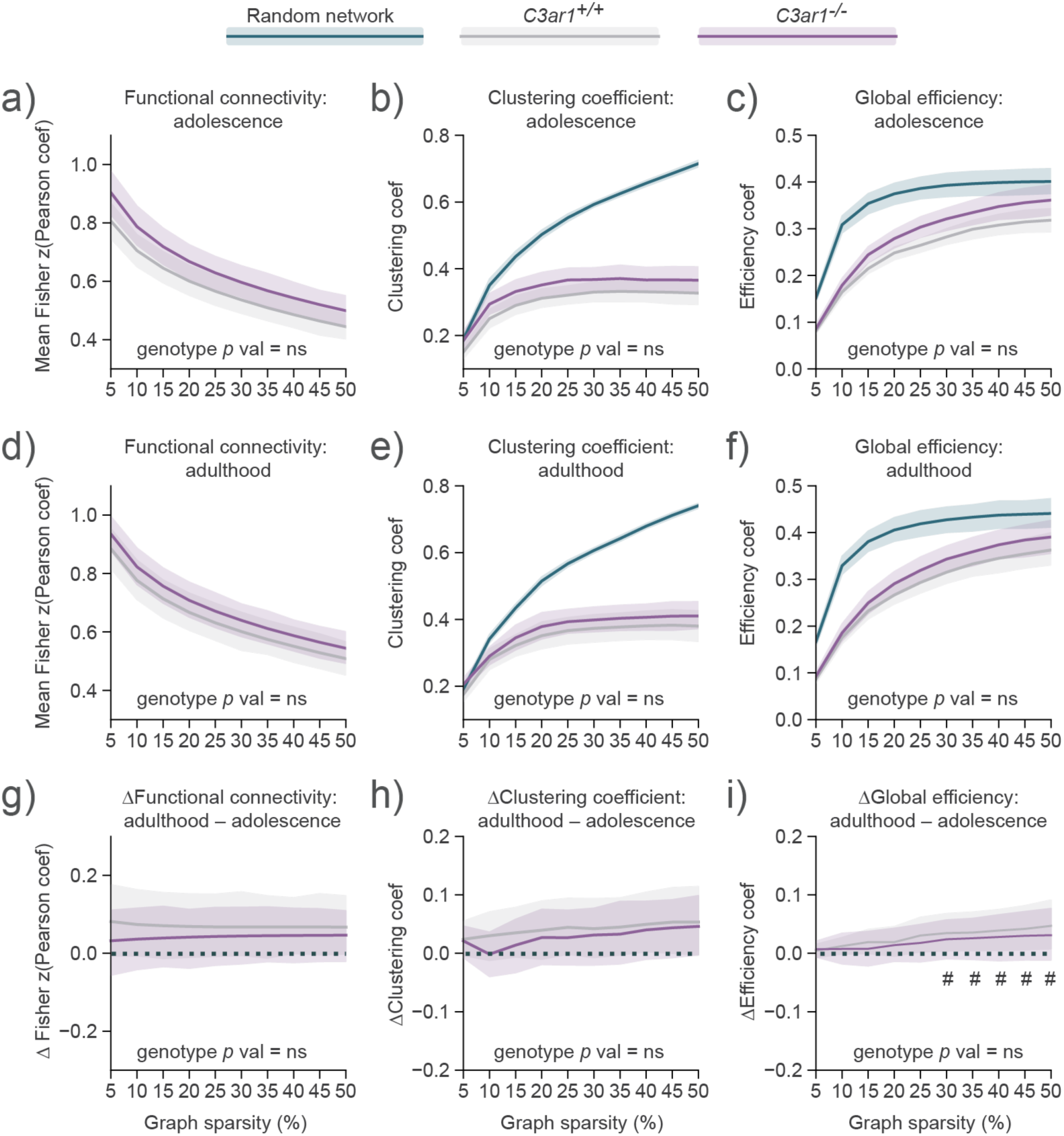
Global functional connectivity is not changed in *C3ar1*-deficient mice. FC, clustering coefficient and global efficiency at decreasing graph sparsity levels in adolescence **(a–c)**, and adulthood **(d–f)**, and change over time **(g–i)**. **(a–i)** Statistical significance was determined with sexes combined using AUC permutation testing (10,000 iterations) and the resulting *p* values were corrected for multiplicity with the BH method (n = 3 tests per outcome measure). Adolescence: *C3ar1^+/+^* n = 32 (15 males, 17 females); *C3ar1^-/-^* n = 32 (18 males, 14 females); Adulthood: *C3ar1^+/+^* n = 33 (16 males, 17 females) and *C3ar1^-/-^* n = 32 (17 males, 15 females); change: *C3ar1^+/+^* n = 30 (14 males and 16 females); *C3ar1^-/-^* n = 31 (17 males and 14 females). Data are shown as mean ± 95% CI. **(g–i)** Two-sided one-sample *t* tests (difference from 0) for global connectivity changes at each sparsity level with genotypes combined, corrected using the BH procedure (n = 10 sparsity levels), # = adjusted *p* value < 0.05.

We applied graph theory to characterise global brain network connectivity across all regions, focusing on two key metrics: clustering coefficient (Figure 4b, **e** and **h**) and global efficiency (Figure 4c, **f** and **i**), and applying the same strategy for proportional thresholding as for FC. The clustering coefficient reflects the tendency of nodes to form connected local clusters, with higher values indicating the presence of more highly interconnected subnetworks within the brain (Bullmore and Sporns 2009).

Global efficiency refers to the average of shortest paths linking nodes in a network and can be a proxy of information integration abilities since it decreases with cognitive deficit (Berlot et al. 2016; Hawkins et al. 2020) and increases with development (Jiang et al. 2023). Clustering coefficient and global efficiency appeared higher in *C3ar1*-deficient animals at both time-points, but this was not significant (**Supplemental table 1)**. Further, we found no effects of genotype across global connectivity measures when we treated males and females as separate groups (**Table 1**, for *p* values, see **Supplemental table 2**).

**Table 1.** Mean network connectivity metrics ± 95% CI in males and females of both genotypes.

| Age | Adolescence |  |  |  | Adulthood |  |  |  |
| --- | --- | --- | --- | --- | --- | --- | --- | --- |
| *C3ar1* status | +/+ | -/- | +/+ | -/- | +/+ | -/- | +/+ | -/- |
| Sex | F | F | M | M | F | F | M | M |
| Connectivity metric (AUC ± 95% CI) | | | | | | | | |
| FC | 25.5<br>± 2 | 27.5<br>± 3.1 | 25.9<br>± 5 | 29.5<br>± 4.5 | 27.6<br>± 3.3 | 28.9<br>± 2.9 | 29.9<br>± 5.3 | 31.9<br>± 4.7 |
| GE | 11.5<br>± 0.8 | 12.7<br>± 1.2 | 11.4<br>± 1.5 | 13.2<br>± 1.7 | 12.17<br>± 1.2 | 12.9<br>± 1.3 | 13.3<br>± 2 | 14.5<br>± 2.1 |
| CC | 13.5<br>± 1.1 | 14.9<br>± 1.9 | 13.9<br>± 2.8 | 15.9<br>± 2.6 | 14.8<br>± 2 | 15.8<br>± 1.9 | 16.3<br>± 3.2 | 17.4<br>± 2.6 |

FC, global efficiency and clustering coefficient appeared to increase with brain maturation when genotype groups were combined, but this was only significant in the case of global efficiency when graph sparsity was lower, that is, when 30–50% of the strongest connections were retained, with small effect sizes observed (Cohen’s *d* = 0.30–0.32). These findings indicate that while *C3ar1* deficiency did not result in alterations in global network properties under our experimental conditions, developmental changes in global efficiency were detectable.

### *C3ar1*-deficiency does not influence functional brain networks

Since *C3ar1*-deficient mice did not show statistically significant changes in global connectivity metrics, we next examined specific networks after conducting *t*-tests for each pairwise ROI FC between genotypes. For this we used two hypothesis-free approaches; false discovery rate (FDR) correction to identify strongly differing edges between genotypes, and network-based statistics (NBS) (Zalesky, Fornito, and Bullmore 2010) to detect network-level differences while controlling for family-wise error rate. Thresholding the resulting *t*-value matrices at |*t*| ≥ 2 for adulthood and adolescence, and their changes over time (adulthood – adolescence; Figure 5a), no edges remained significant after FDR correction. Then, using NBS, which tests the likelihood of detecting a connected component of a specific size, we determined that the network sizes observed after thresholding at |*t*| ≥ 2 (n = 68 at adolescence, n = 15 at adulthood, and n = 1 for change) could occur by chance in these datasets.

**Figure 5.**
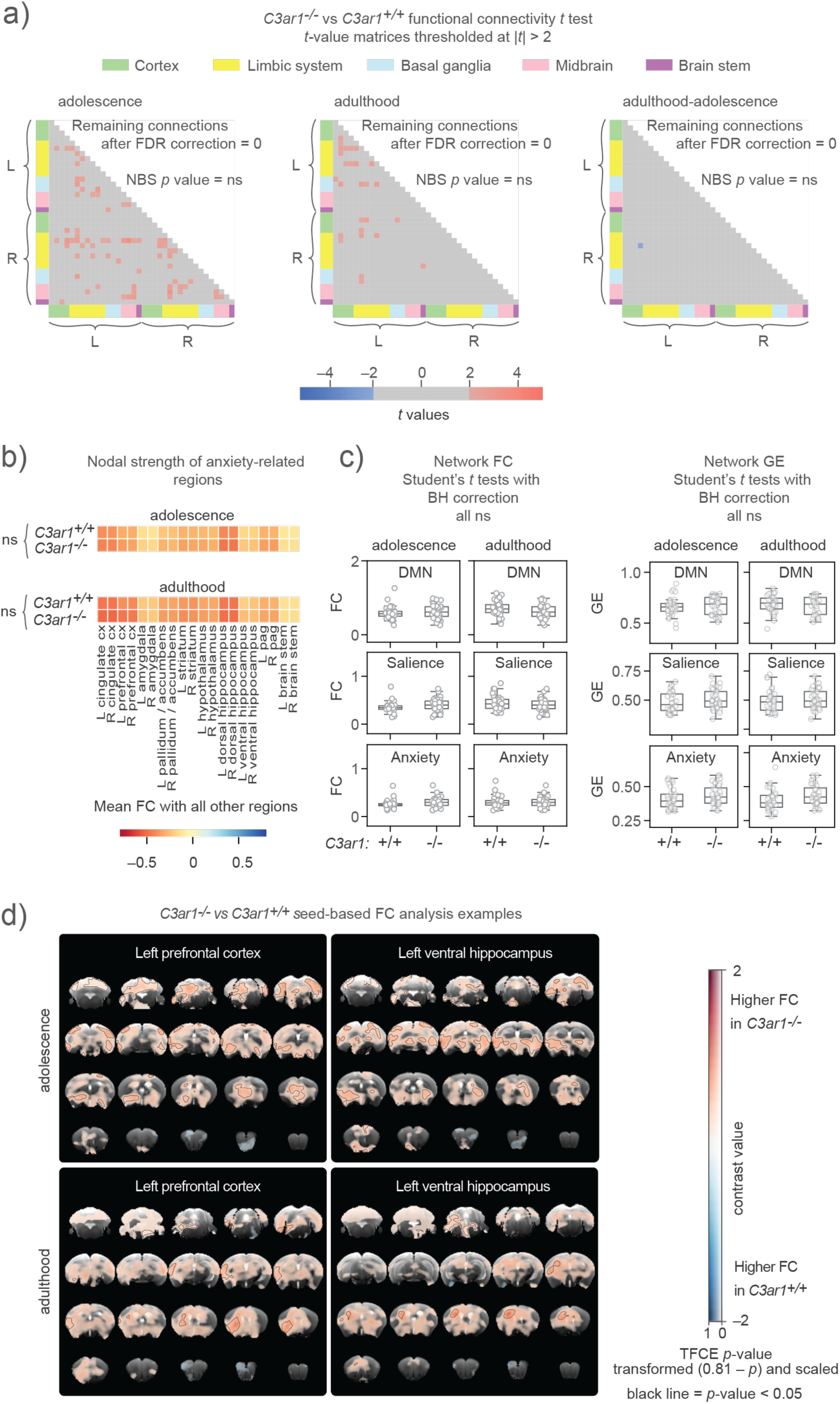
*C3ar1*-deficiency does not influence functional brain networks. **(a)** Thresholded adjacency matrices comparing *C3ar1^+/+^* vs *C3ar1^-/-^* groups where |*t*| ≥ 2 in adolescence and adulthood, and the change between time points. No connections with |*t*| ≥ 2 remained significant after the false discovery rate (FDR) correction. Additionally, the number of connections in the |*t*| ≥ 2 adjacency matrix components were not significant when assessed using network-based statistics (NBS) correction. **(b)** Nodal strength (mean absolute connectivity) of *a priori* anxiety-related seeds (e.g. L cingulate cortex correlation coefficients with all other regions). *P* values were calculated using a mixed ANOVA with between-subjects factor of genotype and within-subjects factor of region. Only the region effect was significant at both time-points (both *p* values < 0.001, η_p_^2^ = 0.69 in adolescence and η_p_^2^ = 0.73 in adulthood). **(c)** Mean functional connectivity (FC) and global efficiency (GE). Student’s *t* tests within a metric (n = 4) corrected for multiple comparisons using the BH method. DMN = default mode network **(d)** Examples of seed-based FC maps showing voxel-wise group differences between *C3ar1^-/-^* and *C3ar1^+/+^* mice (using *t* tests) with seeds placed in the left ventral hippocampus and left prefrontal cortex in adolescence and adulthood datasets. The dual scale bar displays contrast value on the x-axis and TFCE *p*-values (transformed 0.81 – p) on the y-axis. The transformed *p*-values have been re-scaled to range from 0 to 1 for visualisation, with darker colours representing greater statistical significance. Black outlines demarcate regions where TFCE *p*-values are below 0.05.

Given that neither one of the hypothesis-free approaches, FDR correction and NBS, detected any genotype-related differences, we next focused on anxiety-related regions. This decision was motivated by prior evidence of anxiety-like behaviour in *C3ar1*-deficient animals (Westacott et al. 2022) and the inclusion of anxiety-specific tests in our behavioural battery. We calculated the mean FC, or nodal strength, of 20 *a priori* selected brain regions relevant to anxiety. We did not detect effects of genotype (Figure 5b), sex (**Table 2**) or sex-by-genotype interaction (**Supplemental table 3**).

**Table 2.** Mean anxiety-associated node connectivity in males and females of both genotypes.

| Age | Adolescence |  |  |  | Adulthood |  |  |  |
| --- | --- | --- | --- | --- | --- | --- | --- | --- |
| *C3ar1* status | +/+ | -/- | +/+ | -/- | +/+ | -/- | +/+ | -/- |
| Sex | F | F | M | M | F | F | M | M |
| Brain area (mean node connectivity strength) |  |  |  |  |  |  |  |  |
| L cingulate cx | 0.39 | 0.41 | 0.39 | 0.44 | 0.45 | 0.49 | 0.48 | 0.54 |
| R cingulate cx | 0.38 | 0.41 | 0.38 | 0.43 | 0.45 | 0.49 | 0.48 | 0.53 |
| L prefrontal cx | 0.29 | 0.33 | 0.32 | 0.37 | 0.41 | 0.44 | 0.37 | 0.49 |
| R prefrontal cx | 0.28 | 0.28 | 0.33 | 0.37 | 0.41 | 0.45 | 0.39 | 0.47 |
| L amygdala | 0.14 | 0.17 | 0.15 | 0.2 | 0.2 | 0.2 | 0.24 | 0.28 |
| R amygdala | 0.11 | 0.16 | 0.15 | 0.18 | 0.19 | 0.2 | 0.25 | 0.25 |
| L pallidum & accumbens | 0.23 | 0.27 | 0.23 | 0.31 | 0.3 | 0.29 | 0.35 | 0.4 |
| R pallidum & accumbens | 0.24 | 0.26 | 0.24 | 0.3 | 0.31 | 0.31 | 0.35 | 0.39 |
| L striatum | 0.28 | 0.32 | 0.3 | 0.36 | 0.32 | 0.36 | 0.4 | 0.43 |
| R striatum | 0.26 | 0.31 | 0.32 | 0.35 | 0.33 | 0.36 | 0.42 | 0.41 |
| L hypothalamus | 0.28 | 0.29 | 0.27 | 0.34 | 0.32 | 0.32 | 0.34 | 0.38 |
| R hypothalamus | 0.25 | 0.27 | 0.26 | 0.32 | 0.31 | 0.32 | 0.33 | 0.37 |
| L dorsal hippocampus | 0.4 | 0.43 | 0.42 | 0.49 | 0.44 | 0.46 | 0.48 | 0.52 |
| R dorsal hippocampus | 0.41 | 0.43 | 0.42 | 0.49 | 0.44 | 0.47 | 0.5 | 0.52 |
| L ventral hippocampus | 0.18 | 0.18 | 0.17 | 0.26 | 0.21 | 0.23 | 0.2 | 0.27 |
| R ventral hippocampus | 0.18 | 0.23 | 0.19 | 0.25 | 0.22 | 0.25 | 0.25 | 0.26 |
| L PAG | 0.28 | 0.3 | 0.3 | 0.38 | 0.27 | 0.3 | 0.32 | 0.36 |
| R PAG | 0.28 | 0.31 | 0.3 | 0.39 | 0.27 | 0.29 | 0.32 | 0.37 |
| L brain stem | 0.11 | 0.11 | 0.12 | 0.18 | 0.1 | 0.14 | 0.16 | 0.16 |
| R brain stem | 0.11 | 0.09 | 0.12 | 0.18 | 0.11 | 0.13 | 0.17 | 0.18 |

Next, we examined FC and global efficiency within two resting state networks that have been linked to anxiety and emotionality in humans (Coutinho et al. 2016; Geng et al. 2015; Schimmelpfennig et al. 2023) and that have been observed in mice (Grandjean et al. 2020; Sforazzini et al. 2014; Hikishima et al. 2023); the default mode network (DMN) and the salience network (Figure 5c). Additionally, we analysed network connectivity within a third, anxiety network (regions in Figure 5b). Consistent with our earlier findings, we did not observe genotype-dependent differences in these networks at either time-point.

The only significant genotype-related differences were seen in voxel-wise seed-to-brain connectivity analysis. Two anxiety-related regions, the left prefrontal cortex and left ventral hippocampus (Figure 5d; **Supplemental** figure 8 for the right hemisphere), had a weak but significant and widespread higher connectivity in *C3ar1*-deficient animals compared to controls at both time-points. Seed-based analysis further revealed higher FC across the brain when other anxiety-related regions were used as seeds, mostly in adolescence (**Supplemental** figure 9).

However, this effect was not confined to anxiety-related regions or specific networks (**Supplemental** figure 10). Although the increase of FC in *C3ar1*-deficient mice appears widespread, this is not robust, since our seed-based analysis controls for voxel-wise comparisons within subjects but does not correct for testing multiple seeds.

### *C3ar1*-deficient mice do not have discernible behavioural phenotypes

Behavioural testing is another way of assessing functional consequences of genetic manipulations with expected neurodevelopmental sequelae (Crawley 2007). We aimed to evaluate the impact of *C3ar1* deficiency on anxiety-like behaviour, locomotion and recognition memory, which were chosen based on prior reports of abnormalities in *C3ar1*-deficient mice. We used a battery of well-established behavioural tests, including the OF test and EPM for anxiety-like behaviour and locomotion, as well as NOR for recognition memory. We also tested PPI which is a sensorimotor reflex consistently found to be attenuated in schizophrenia (Ludewig, Geyer, and Vollenweider 2003; Mena et al. 2016) but which has hitherto not been tested in *C3ar1*-deficient mice before.

We performed behavioural testing in both cohorts described in the structural MRI results sections. In Cohort 1 (Figure 2a), behavioural testing (EPM, then OF) occurred shortly before the adulthood MRI scan (and two months after adolescence MRI). Behaviour of Cohort 2 was also tested in adulthood (OF, NOR, EPM, PPI, in that order) followed by MRI, except that these mice were not previously scanned under anaesthesia. Behavioural data from each cohort were analysed separately to account for differences in study design (see also Methods).

To assess multiple measures of anxiety-like behaviour and locomotion along one another (Figure 6a**–b****, Supplemental table 4** for description of anxiety-like metric selection), we calculated z-scores for each behavioural outcome measure for *C3ar1*-deficient mice relative to wild-type controls (**Table 3** for untransformed means). In Cohort 1, no significant genotype effects were detected in anxiety-like behaviour or locomotion at either time-point (**Supplemental table 5**). To rule out any potential confounding effects from co-housing littermate mutants and wild-types (Kalbassi et al. 2017), we also examined whether the number of *C3ar1*-deficient cage-mates influenced wild-type behaviour but found no consistent pattern or evidence of systematic anxiety-like effects in wild-types (**Supplemental** figure 11).

**Figure 6.**
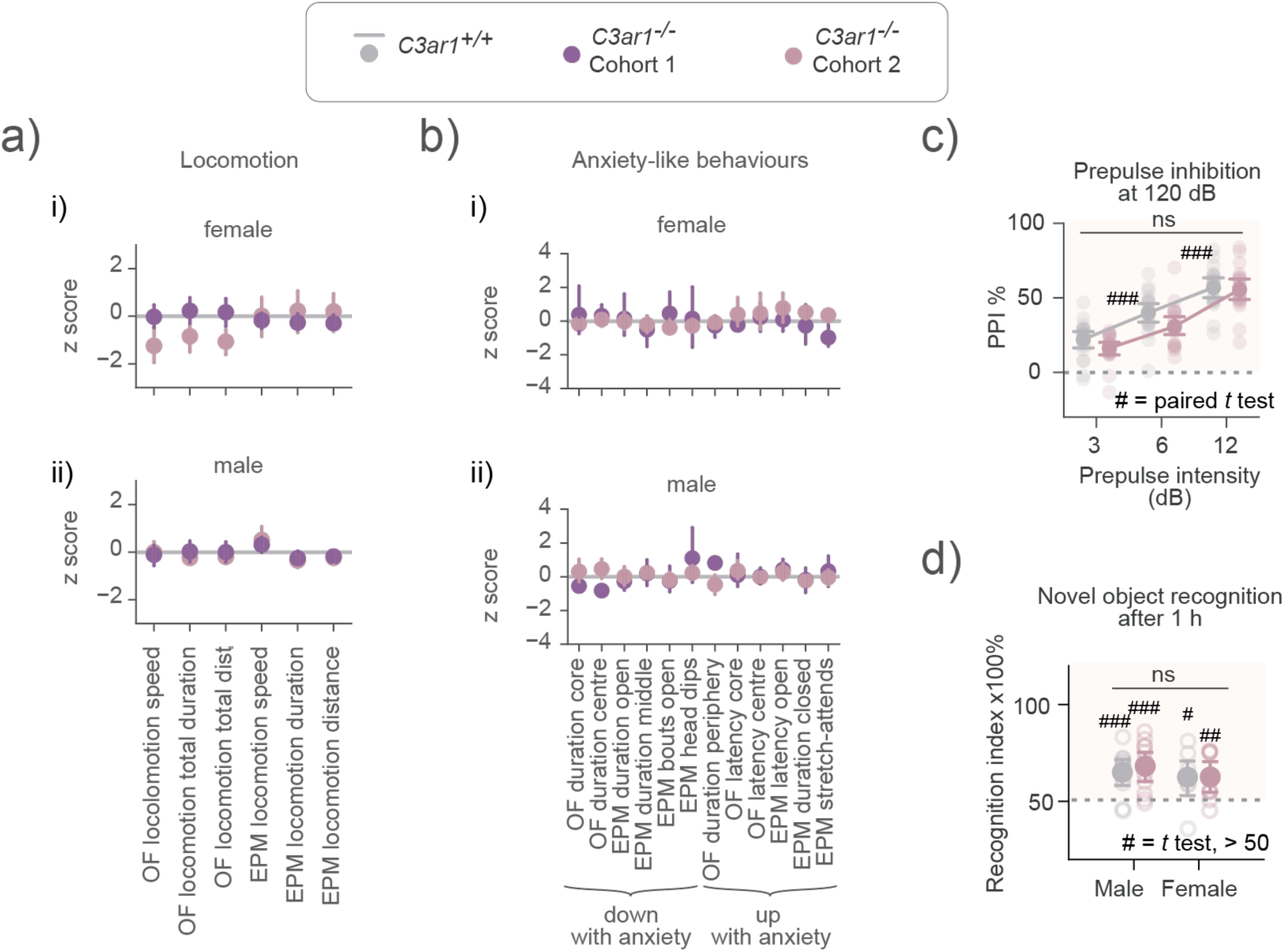
*C3ar1* deficiency does not cause behavioural abnormalities. **(a)** Z-scored (normalised to *C3ar1^+/+^* control mean = 0) locomotion metrics for *C3ar1^-/-^* animals from EPM and OF tests, showing results from Cohort 1 and 2. Two-way ANOVA for Cohort 1 (4 groups, males and females separated) or Student’s *t* tests for Cohort 2 (2 groups, males and females combined due to smaller sample size) where parametric assumptions were met, otherwise the Kruskal Wallis test (with Dunn in Cohort 1); uncorrected for multiplicity, all ns. **(b)** Same as (a) but for anxiety-like metrics from EPM and OF tests; *p* values all ns. **(c)** Prepulse inhibition in Cohort 2. PPI increased with increasing prepulse intensity in both, *C3ar1^+/+^* vs *C3ar1^-/-^* mice (paired *t* tests p value ### < 0.001). Dotted line at 0 indicates inhibition threshold. A score above 0 indicates inhibition (shaded area). Slope plot shows means and 95% confidence intervals. Individual mice are plotted three times at increasing prepulse intensity. Prepulse inhibition did not differ by genotype at any prepulse intensity (uncorrected Student’s *t* tests at 3 dB, 6 dB and 12 dB, *t*_[37]_ = 1.58, 1.99, 0.28, *p* = 0.12, 0.54, 0.17). **(d)** Novel object recognition (NOR) recall 1 hr after acquisition in Cohort 2. Both groups showed novelty preference (one-sample *t* test, value > 50/chance, # = *p* < 0.05, ## = *p* < 0.01 ### = *p* < 0.001). There were no differences between groups (Student’s *t* test *t*_[37]_ = 0.42, *p* = 0.67). The dotted line marks the chance threshold. **(a-d)** Cohort 1: *C3ar1^-/-^* n = 33 (17 male, 16 female), *C3ar1^+/+^* n = 33 (16 male, 17 female), Cohort 2: *C3ar1^-/-^* n = 20, *C3ar1^+/+^* n = 19 (males and females combined). Data are expressed as mean ± 95% CI.

**Table 3.**
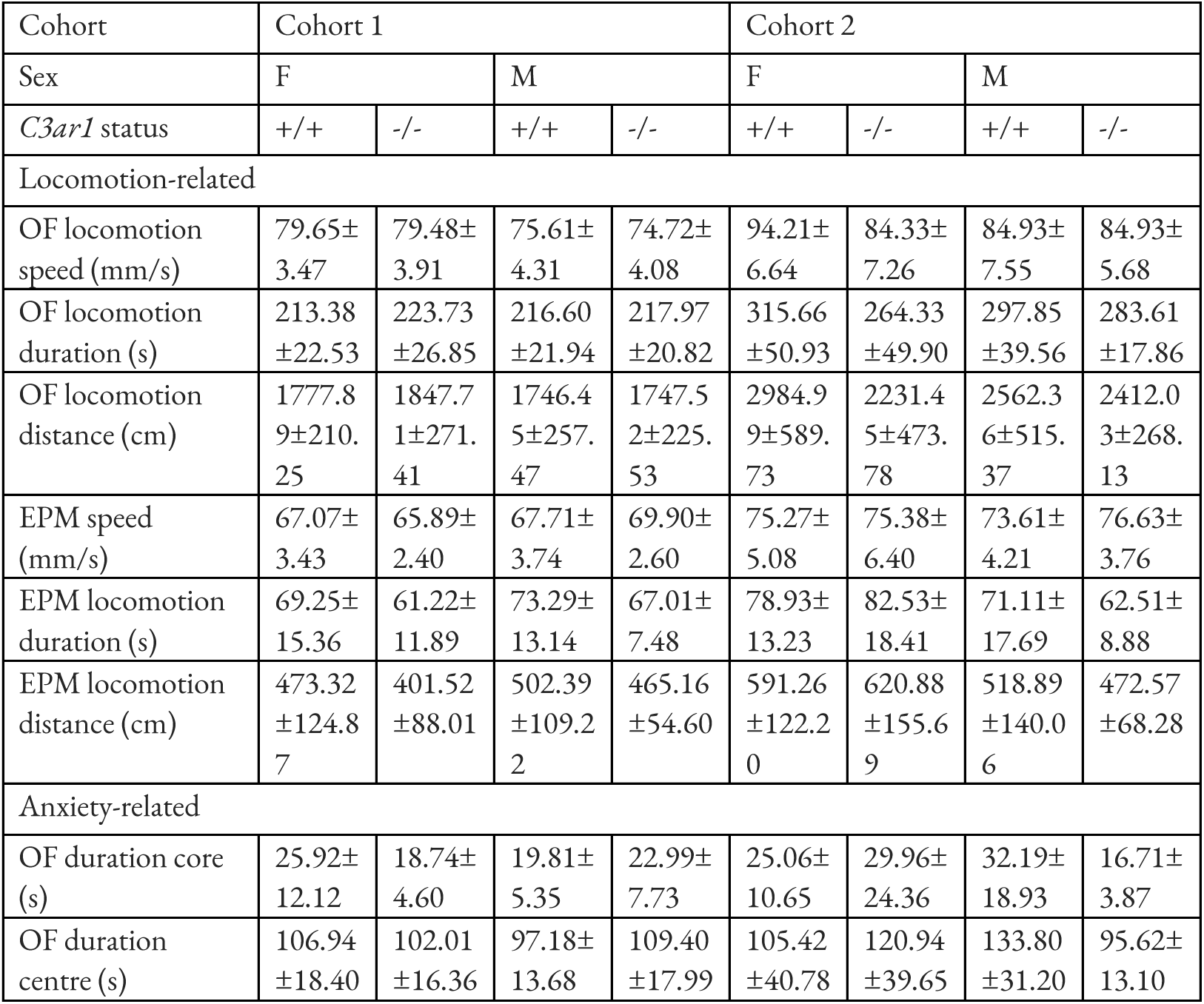
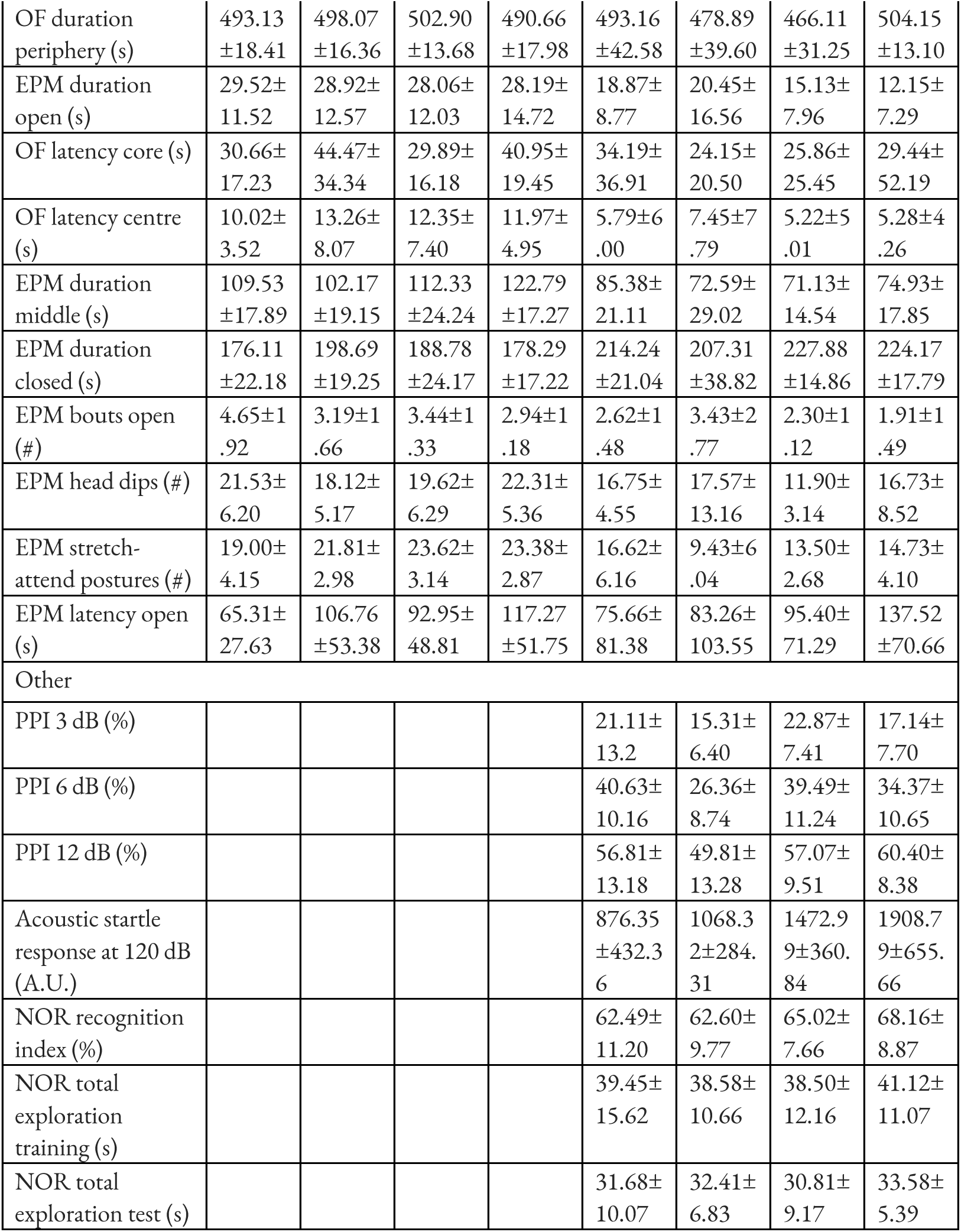
Mean behavioural outcome measures ± 95% CI in males and females of both genotypes.

In Cohort 2, most anxiety-related and locomotion measures showed no genotype differences (**Supplemental table 6**), except for reduced distance travelled in the centre of the OF arena by *C3ar1*-deficient mice (uncorrected two-sample *t*-test *p* < 0.01, Cohen’s *d* = 0.92, **Supplemental table 6**). However, no genotype effect on distance was detected in the core of the OF arena or in the centre and core of the OF arena in Cohort 1.

The sample size (n = 38) of Cohort 2 was too small to reliably test for interactions between sex and genotype, meaning we only had enough statistical power to detect very large effects (Cohen’s *f* = 0.5 at alpha = 0.05 and 80% power), which were clearly not observed across behavioural measures. No sex-by-genotype interactions were observed in Cohort 1 (**Supplemental table 5**).

Additionally, PPI testing (performed exclusively on Cohort 2) showed no effect of *C3ar1* deletion, though PPI increased with prepulse intensity as expected (Figure 6c), confirming that the test was set up to reliably measure this phenomenon. NOR testing (also performed exclusively on Cohort 2) revealed no genotype differences, with all groups demonstrating successful learning based on recognition indices significantly above chance levels (Figure 6d). Overall, these findings suggest that *C3ar1* deficiency does not result in robust anxiety-like or hyperactive phenotypes, nor deficits in recognition memory or PPI.

**Table 4.**
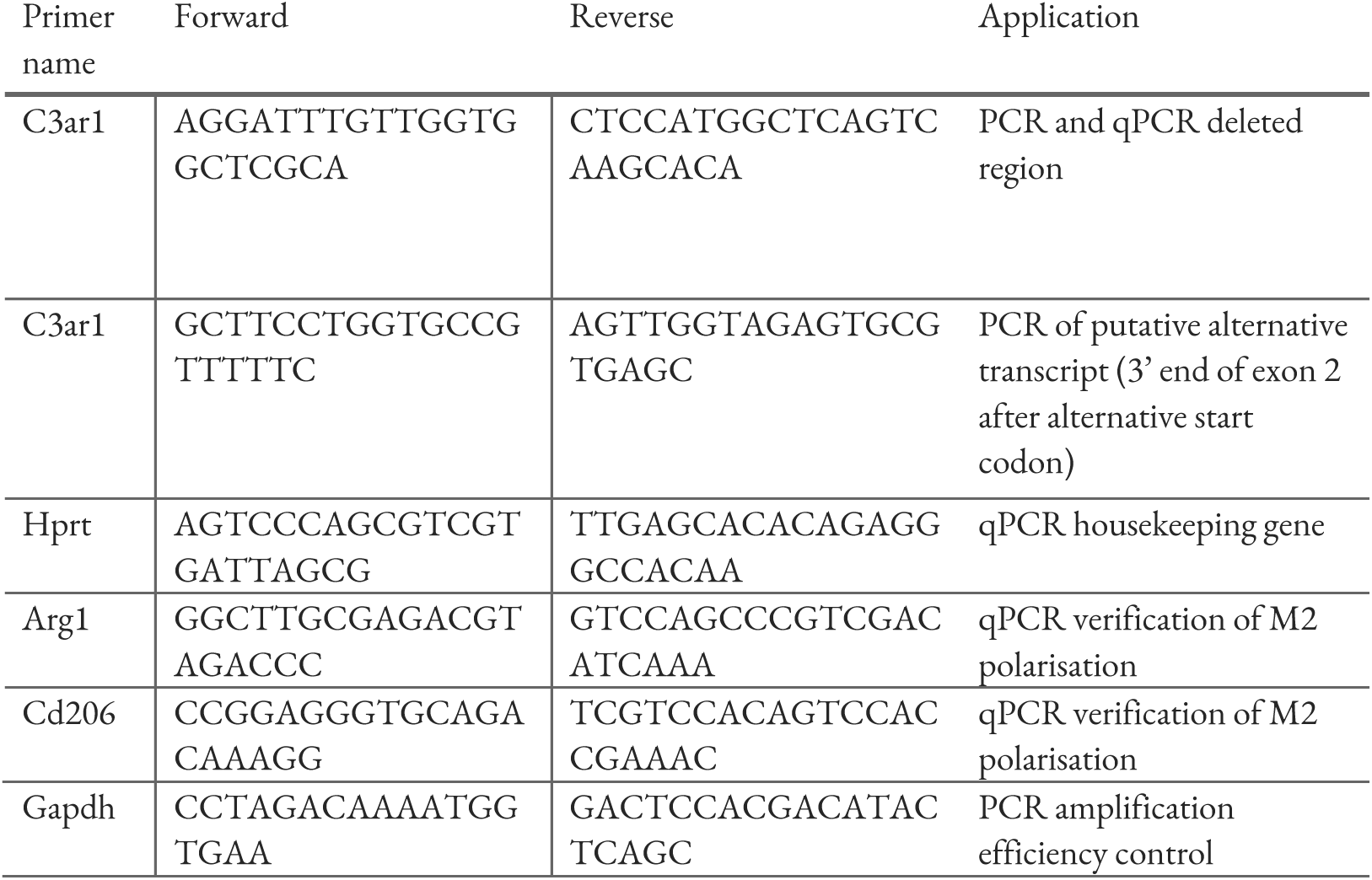
Primer sequences.

**Table 5.** PCR reaction.

| Amount (µl) | Reagent |
| --- | --- |
| 12.5 | GoTaq® G2 Master Mix (Green) |
| 1 | *C3ar1* forward primer, 10 mM |
| 1 | *C3ar1* reverse primer, 10 mM |
| 1 | *Gapdh* forward primer, 10 mM |
| 1 | *Gapdh* reverse primer, 10 mM |
| 7.5 | H<sub>2</sub>O |
| 1 | Template DNA |
| Final volume: 25 µl |  |

**Table 6.** Cycling conditions.

| Step | Time | Temperature |
| --- | --- | --- |
| 1. Initial denaturation | 5 min | 94°C |
| 2. Denaturation | 30 sec | 94°C |
| 3. Annealing | 30 sec | 60°C |
| 4. Extension | 1 min | 72°C |
| 5. Repeat steps 2-4 | Repeat 35 x |  |
| 6. Final extension | 5 min | 72°C |

**Table 7.** MRI acquisition parameters.

| Image type | TR (ms) | TE (ms) | Flip angle (°) | Averages | Bandwidth (kHz) | FOV | Matrix | Other |
| --- | --- | --- | --- | --- | --- | --- | --- | --- |
| AFI | 20/<br>100 | 2.85 | 55 | 1 | 25 | 16.2 x<br>16.2 x 9 | 42 x 42 x<br>24 |  |
| MTw | 24 | 2.5 | 6 | 2 | 100 | 16.2 x<br>16.2 x 9 | 108 x<br>108 x 60 | 6 echoes with 2.1 ms spacing;<br>MT pulse: gaussian, 4 ms, amplitude 10 μT, offset -3 kHz, bandwidth 685 Hz |
| PDw | 20 | 2.5 | 4 | 2 | 100 | 16.2 x<br>16.2 x 9 | 108 x<br>108 x 60 | 7 echoes with 2.1 ms spacing |
| T1w | 20 | 2.5 | 20 | 2 | 100 | 16.2 x<br>16.2 x 9 | 108 x<br>108 x 60 | 7 echoes with 2.1 ms spacing |
| T2w<br>in vivo | 5000 | 42 | 90/<br>180 | 4 | 50 | 16 x 12 | 128 x 96 | 32 slices, slice thickness 0.5, RARE factor 8 |
| T2w<br>ex vivo | 300 | 30 | 90/<br>180 | 1 | 50 | 25 x 24<br>x 18 | 250 x<br>240 x<br>180 | RARE factor 4, scan time = 1 h 5 min |
| DWI<br>in vivo | 3000 | 21 | 90/<br>180 | 2 | 357 | 19.2 x 12 | 96 x 60 | Single shot spin-echo planar imaging, 30 slices, slice thickness 0.5 mm, three diffusion shells: $b = 350 \text{ s/mm}^2$ with 9 directions, $b = 1000 \text{ s/mm}^2$ with 34 directions, $b = 2000 \text{ s/mm}^2$ with 78 directions ($\delta = 3 \text{ ms}$, $\Delta = 11 \text{ ms}$), 3 $b_0$ images per shell |
| DWI<br>ex vivo | 300 | 28.5 | 90/<br>180 | | 300 | 25 x 24<br>x 18 | 200 x<br>192 x<br>144 | Stejskal-Tanner pulsed gradient spin echo sequences with a 3D segmented EPI readout. 12 segments and a total of 90 diffusion-weighted images acquired at a $b$-value of $4000 \text{ s/mm}^2$ ($\delta = 4 \text{ ms}$, $\Delta = 13 \text{ ms}$) |
| BOLD<br>fMRI | 1000 | 15 | 55 | 1 | 200 | 20 x 20 | 64 x 64 | Single-shot gradient-echo EPI sequence, 16 slices, slice thickness 0.5, 720 repetitions |

## Discussion

Here we used longitudinal neuroimaging and behavioural testing to investigate the impacts of *C3ar1* deletion on brain structure and function across early life development in mice. We found no robust evidence that *C3ar1* deficiency affects total or regional brain volume, white matter FA, global FC, global efficiency, clustering coefficient, or specific functional networks.

We also found no altered behavioural phenotype in either male or female *C3ar1*-deficient mice in adulthood. These results raise questions regarding the source of discrepancies between these data, our previous behavioural work (Westacott et al. 2022; Westacott et al. 2021), and that of others (Crider et al. 2018; Coulthard et al. 2018; Pozo-Rodrigálvarez et al. 2021; Sun et al. 2024), which may be attributed to our use of littermate controls in this study, as will be elaborated upon in the following sections. Overall, the lack of a behavioural phenotype combined with the paucity of genotype effects on brain structure across time-points means that the role of C3aR1 in neural development and behaviour is not as significant as originally thought, at least in the absence of exacerbating environmental or immune triggers.

### Absence of *C3ar1*-dependent effects on brain structure

Our morphometric analysis showed that total and regional brain volumes were not changed between *C3ar1*-deficient and wild-type mice. We were, however, able to detect sex-specific neurodevelopmental changes that have previously been reported in rodents, such as brain growth occurring later in females, minimal total and regional brain volume differences in adulthood and larger volume of MEPO and BNST in males (Qiu et al. 2018; Guma et al. 2024; Gorski et al. 1978; Hines, Allen, and Gorski 1992; de Courten-Myers 1999), supporting the sensitivity of our methodology. Together with the low variability in our sample, our ability to detect these known sex differences means that we were well-positioned to identify potential genotype effects had they been present.

### Absence of *C3ar1*-dependent effects on white matter fractional anisotropy

*C3ar1* is predominantly expressed by microglia in the brain (Quell et al. 2017; Tasic et al. 2018), where it influences their reactivity and phagocytosis (Vasek et al. 2016; Gedam et al. 2023; Zheng et al. 2021; Lian et al. 2015; Gnanaguru et al. 2023).

Notably, changes in microglial characteristics have been linked to developmental alterations in FA in both mice and humans (Falangola et al. 2023; Chan et al. 2024). We found no *C3ar1*-dependent changes in the white matter despite showing the overall effect of time on the brain maturation on FA, which is known to increase during development between adolescence and adulthood in both humans and mice (Brouwer et al. 2012; Hagmann et al. 2010; Reynolds et al. 2019; Piekarski et al. 2023; Chahboune et al. 2007). Lack of a phenotype after the deletion of a predominantly microglial gene may not be surprising since mice that completely lack microglia for their entire lifespan (achieved by deleting the super enhancer for macrophage colony-stimulating factor receptor, *Csf1r*) do not have overt neurodevelopmental phenotypes (Rojo et al. 2019), and only show vulnerability in a pathological context involving neuroinflammation (Kiani Shabestari et al. 2022). This raises the possibility that neurodevelopmental phenotypes associated with *C3ar1* deficiency may likewise only become apparent under neuroinflammatory conditions.

### Widespread, weak seed-to-brain functional connectivity increase in *C3ar1*-deficient mice in adolescence

While we did not detect genotype-dependent global FC changes nor changes to clustering coefficient and global efficiency at either time-point, we did observe a genotype-independent developmental increase in global efficiency with time, which has been reported before (Jiang et al. 2023; Hagmann et al. 2010; Koenis et al. 2018), particularly at lower sparsities (Cai, Dong, and Niu 2018). This additionally suggests that our method was sensitive enough to pick up developmentally relevant effects.

Similarly to whole brain analyses, we did not observe any genotype-dependent changes to specific brain networks, including anxiety networks (Figure 5), which we hypothesised to be affected based on our previous behavioural results (Westacott et al. 2022). However, our anxiety network connectivity null findings are internally consistent with the absence of an anxiety-like phenotype in the cohorts tested in this study. Combined with a recent null report for an anxiety-like effect in the EPM in *C3ar1*-deficient animals by another group (Sun et al. 2024), our data indicate that the effect of *C3ar1* on anxiety-networks and anxiety-like behaviour is context-dependent if present.

Although we did not detect network-specific effects, *C3ar1*-deficient mice exhibited subtle but widespread increases in resting-state FC in voxel-wise seed-based analyses during adolescence across nearly all seeds examined, with a weaker effect persisting into adulthood (Figure 5 and **Supplemental** figures 8-10). These increases in connectivity may reflect developmental alterations in circuit properties caused by the absence of C3aR1. Similar findings have been reported for other microglial receptors—such as TREM2, CX3CR1, and CR3—where genetic deficiency altered adult brain FC (Filipello et al. 2018; Zhan et al. 2014; Deivasigamani et al. 2023). In these reports, TREM2 or CX3CR1 knockouts led to impaired synapse elimination alongside decreased FC, accompanied by social behaviour deficits and increased repetitive behaviour (Filipello et al. 2018; Zhan et al. 2014). In contrast, CR3 knockout mice had no deficits in synapse or axon refinement but showed reduced phagocytosis of perinatal cortical neurons and higher cortical FC (Deivasigamani et al. 2023). It is indeed possible that similarly to CR3, C3aR1 is also involved in perinatal neuron phagocytosis so that its deletion results in higher number of neurons in early embryonic development and a weak brain-wide increase in FC, particularly as *C3ar1*-deficiency leads to a deficit of developmental astrocyte phagocytosis in the retina (Gnanaguru et al. 2023). For now, we urge caution with this interpretation for two reasons that should be considered together. These connectivity changes were isolated findings–that is, they were not accompanied by any structural alterations, which would be unexpected if a larger number of neurons were present, nor were they associated with any behavioural changes. Second, the voxel-wise seed-based analysis is the least robust presented here, as the observed changes were largely transient, weak, not confined to a specific subnetwork, and not corrected for multiple comparisons across seeds. Future studies investigating FC in *C3ar1*-deficient mice should incorporate repeated measures from the same individual during adolescence to improve robustness as well as assessing synaptic transmission more directly, for example by electrophysiological assessment in brain slices, although it is still unclear which brain regions should be targeted currently with the latter approach.

### Behavioural outcomes were unaffected by *C3ar1* deficiency

Neither anxiety-like behaviours nor locomotor activity showed significant differences between genotypes across multiple testing paradigms, apart from a small decrease in ambulation of *C3ar1*-deficient mice in the aversive central zone of the OF arena, which in the absence of a locomotor phenotype could index anxiety-like behaviour (Prut and Belzung 2003). This latter change, however, was not reproducible across study cohorts and was no longer observed in the same study in the core of the OF arena. Tests for recognition memory and sensorimotor gating, NOR and PPI tests, also yielded null results.

Differences between our current findings and previous reports of anxiety-like behaviours (Westacott et al. 2022), hyperactivity (Pozo-Rodrigálvarez et al. 2021) and cognitive deficits (Coulthard et al. 2018) may stem from variations in experimental design, particularly environmental factors such as lighting and testing time. Our behavioural experiments were conducted during the active phase (7 PM– 11 PM, lights off) to minimise stress from sleep deprivation, which is known to upregulate complement in the brain (Tillmon et al. 2024; Crider et al. 2018; Li et al. 2023; Tripathi et al. 2021). It is possible that previous studies, which tested animals during their inactive phase, introduced a stress-dependent “second hit” to *C3ar1* deficiency. However, this seems unlikely since inhibiting complement, including *C3ar1*, has been shown to improve resilience to chronic stress by preventing anhedonia-like behaviour and neuroinflammation (Li et al. 2023; Crider et al. 2018; Tripathi et al. 2021; Madeshiya et al. 2022; Tillmon et al. 2024).

Unlike previous studies of *C3ar1*-deficient mice, our study employed littermate control animals, meaning that wild-type mice were co-housed with their *C3ar1*-deficient siblings. This approach minimises potential environmental differences between groups, which is especially important in neurobehavioral studies where subtle environmental factors can significantly affect outcomes. While the use of littermate controls is widely regarded as best practice in such contexts (Holmdahl and Malissen 2012; Valiquette et al. 2023; Bailey, Rustay, and Crawley 2006), it is not without its own potential confounds. For instance, abnormal behaviours exhibited by genetically altered animals can influence the behaviour of co-housed wild-type mice, particularly when the phenotype is pronounced, such as in cases of increased aggression (Kalbassi et al. 2017). However, as aggression has not been reported in *C3ar1*-deficient mice, and their previously described behavioural phenotype was not notably severe, we considered such an influence unlikely in our study, especially as the anxiety-like behaviour of wild-type mice did not seem to depend on the number of *C3ar1*-deficient cage-mates.

The complement system plays an essential role in normal pregnancy and parturition (Girardi et al. 2020). A notable aspect of using a littermate design is that litters of mixed genotype are born from exclusively heterozygous parents, whereas in most prior studies of *C3ar1*-deficiency, both the mother and offspring were homozygous knockouts. This raises the possibility that behavioural effects in *C3ar1*-deficient mice may have resulted from altered intrauterine environments or care by *C3ar1*-deficient mothers. In line with this, the inbred homozygous *C3ar1* knockout colony used in our previous studies showed increased pre-weaning deaths and maternal cannibalism (unpublished data). If this is indeed the reason for the previously observed adult behavioural phenotypes, two scenarios should be considered. In the first scenario, the *C3ar1*-deficient offspring are *uniquely sensitive* to C3aR1-dependent *in utero* conditions or maternal care deficits. In the second scenario, the adult phenotypes are not specific to *C3ar1*-deficiency meaning that wild-type mice would have been similarly affected. These scenarios can be tested by using pup transfer and *in-vitro* fertilisation experiments. For now, we were able to avoid measuring these non-specific effects on behaviour by using a littermate design.

The genetic background of mutants is another factor that varies between laboratories and is known to affect phenotypic expression. For example, heterozygous knockout of the autism-associated gene *CHD8* has varying effects in 33 sub-strains (Tabbaa, Knoll, and Levitt 2023), mirroring heterogeneity observed in human *CHD8*-haploinsufficiency. While that study identified significant variability across sub-strains, these profiles consistently differed from that of wild-type littermate controls. In contrast, our study found no robust genotype-dependent differences in the brain globally and across measures or in behaviour, apart from a single internally non-reproducible behavioural outcome measure, meaning that the effect of C3aR1 on behaviour would have to be entirely dependent on modifying genes or environmental factors.

Finally, genetic drift could have precluded reproducibility of previously reported phenotypes. Mice accumulate spontaneous mutations that rapidly reach homozygosity within small colonies, speeding up drift, a phenomenon recognised as early as the 1980s (Fitch and Atchley 1985). So far, our current work is the first to study *C3ar1* deficiency in behaviour using a littermate design, which minimises the number of genetic loci that differ between the mutant and control mice, all of which could influence measured phenotypes.

### Future directions

Aside from the role of C3aR1 in brain structure and function, many aspects of the basic biology of C3aR1 remain unclear, including its expression pattern in the healthy brain, with little information available on the cell types where it is expressed nor its expression contexts and time-points. It is also unclear which G-proteins C3aR1 signals through in different brain cell types. Future studies could integrate toolkits like TRUPATH, a suite of Gαβγ biosensors for analysing G-protein coupling preferences (Olsen et al. 2020), with transcriptomics to address these questions. This approach would be particularly valuable, as it could enable the use of chemogenetics to activate the same G-protein as C3aR1 in specific cell types and contexts to study its function.

Most importantly, we propose that *C3ar1*-deficiency in development should be studied after an immune challenge. Unlike humans, laboratory mice live in an immune-privileged environment, hence investigating the consequences of a combined immune insult with the genetic deficit would be relevant; or better yet, in the context of *C4A* overexpression, which is a known genetic risk factor for schizophrenia (Sekar et al. 2016) and is known to impact white matter integrity (Caseras et al. 2024).

### Limitations

In this study, we provided high-level global data on *C3ar1*-deficient mice. While our study had notable strengths, including the use of a littermate design and the inclusion of female animals, it is not an exhaustive characterisation of the mutant.

While we measured brain region volumes, we cannot definitively address more reductionist questions, such as cellular composition which would have required single cell RNASeq or antibody staining and microscopy. Similarly, for axonal integrity analysis for which we used the proxy of dMRI-derived FA, the gold standard would have been electron microscopy. However, a more granular approach would have necessitated a trade-off with throughput—something that is hard to justify in the absence of strong hypotheses regarding the developmental expression pattern of *C3ar1*. Additionally, our fMRI analysis relied on a specific parcellation of the brain consisting of 36 regions, and different parcellations could yield varying results (Thirion et al. 2014; Zalesky et al. 2010). We encourage others exploring our datasets to therefore experiment with alternative parcellations.

## Conclusion

Contrary to expectations, we found no evidence for *C3ar1*-dependent effects across imaging measures at either time-point, nor could we replicate previously seen behavioural phenotypes. The absence of detectable phenotypes suggests that C3aR1 plays a minimal role in brain structure and function, at least in the absence of an immune trigger. These findings challenge prior assumptions about its neurodevelopmental significance and necessitate further investigation into C3aR1 function on sensitized backgrounds, such as those involving immune activation.

## Methods and materials

### Animals

All animal procedures complied with the UK Animals and Scientific Procedures Act 1986 and were approved by the local ethical committee at King’s College London (KCL). Homozygous *C3ar1^-/-^* mice were generated by homologous recombination in embryonic stem cells and kindly provided by Dr. Bao Lu and Prof. Craig Gerard (Harvard Medical School, Boston, MA) (Humbles et al. 2000). These mice were subsequently backcrossed onto the C57BL/6J strain for at least 12 generations and maintained on a C57BL/6J background in Professor Wuding Zhou’s laboratory at KCL. For this study, cryopreserved stocks were rederived at KCL and crossed to C57BL/6J mice purchased from Charles River to refresh the genetic background following Jackson’s Laboratories line refreshing protocol.

Experimental animals (*C3ar1^-/-^* and *C3ar1^+/+^* littermates) were generated through heterozygote incrosses and resulting genotypes followed Mendelian ratios. The heterozygote breeders were generated by outcrossing heterozygous mice to bought wild-type Charles River C57BL/6Js. The breeders used to produce the experimental animals were derived either from the first, second or third of these outcrosses. Sibling crosses were not conducted, and parental age was between 2–4 months to minimise genetic drift.

Experimental mice were housed in individually ventilated cages under controlled temperature (20–25°C), humidity (50–60%), and a 12-hour light-dark cycle (lights on at 7:00 AM, lights off at 7:00 PM). Environmental enrichment included nesting materials, tunnels, and chew sticks. Mice had *ad libitum* access to irradiated rodent chow and autoclaved water. Animals were group-housed (2–4 mice per cage), with males and females housed separately after weaning (PND21±2). Genotyping was conducted on ear biopsy DNA by Transnetyx using probes targeting the neomycin cassette for the mutant allele and intron 1 for the wild-type allele. No mismatches were identified through double-genotyping 20% of the study cohorts.

### Validation of mutation

Bone marrow-derived macrophages were obtained from tibias and femurs of eight three-month-old mice (n = 4 *C3ar1^+/+^,* n = 4 *C3ar1^-/-^*) following standard protocols. Bone marrow suspension was filtered through a 40 µm mesh, centrifuged at 450 x g for 5 minutes at 4°C, and treated with NH_4_Cl haemolysis buffer (NH_4_Cl 0.15M, KHCO_3_ 0.01M, EDTA 0.0001M). After a second centrifugation under the same conditions, cells were washed with PBS and resuspended in Gibco RPMI 1640 Medium (Thermo Fisher, #21875-034) supplemented with 50 ng/ml recombinant mouse macrophage colony-stimulating factor (M-CSF, R&D Systems, #416-ML-010/CF), 1% penicillin, 1% streptomycin, and 10% heat-inactivated foetal bovine serum (Sigma, #F9665-50ml).

Cells were seeded at 1×10^6^ cells/ml in six-well plates (six wells per animal) and incubated at 37°C with 5% CO_2_ for 72 hours. On day three, the medium was refreshed, and 50 ng/ml recombinant mouse IL4 (R&D Systems, #404-ML-010/CF) was added to half of the wells to skew them towards M2 phenotype. Incubation continued for an additional 48–72 hours, depending on cell confluence.

RNA was extracted using the ReliaPrep™ miRNA Cell and Tissue Miniprep System (Promega, #Z6211) according to the manufacturer’s instructions. RNA concentration was determined with a NanoDrop spectrophotometer (Thermo Scientific, NanoDrop 2000), yielding values between 25.5 ng/µl and 227.5 ng/µl. Reverse transcription PCR (RT-PCR) was performed using the LunaScript® RT SuperMix Kit (NEB, #E3010L), following the manufacturer’s instructions. Depending on RNA yield, either 100 or 500 ng of RNA was used per reaction.

For cDNA PCR and gel electrophoresis, we used GoTaq® G2 Master Mix (NEB, #M7822). We amplified a 79 basepair (bp) fragment in the deleted region alongside a 372 bp region in the *C3ar1* cDNA that was located outside the deleted region and downstream of an alternative start codon identified through Benchling. *Gapdh* primers were included in each reaction to confirm amplification efficiency. PCR products were analysed on a 2% agarose gel stained with GelRed.

For qPCR, we amplified the previously mentioned 79 bp fragment in the deleted region. Samples were analysed in triplicate on 96-well plates (Applied Biosystems, #4346906) with Luna® Universal qPCR Master Mix (NEB, #M3003L), and readings were obtained using an Applied Biosystems StepOnePlus plate reader. The amplification data were processed with the ^ΔΔ^Ct method, normalising against the housekeeping gene Hypoxanthine phosphoribosyltransferase 1 (*Hprt*). Genotypes were arranged alternately across the plate to minimise bias. To verify M2-like polarisation, the expression of M2-specific markers Arginase 1 (*Arg1*) and the mannose receptor Cluster of differentiation 206 (*Cd206*) were assessed.

### Study design

This study used two separate cohorts of male and female *C3ar1*-deficient and littermate wild-type mice. The main, longitudinal MRI cohort, termed Cohort 1, had *in vivo* MRI performed in adolescence (range 27–31 days) and adulthood (range 81–92 days), and the adulthood MRI scan was preceded by OF and EPM tests. The adolescence time-point was chosen because mice reach puberty approximately between PND24–34 (Semaan and Kauffman 2015; Brust, Schindler, and Lewejohann 2015; Pintér et al. 2007). *Ex vivo* imaging was conducted in Cohort 1’s perfusion-fixed brains after the final adulthood scan.

For Cohort 2, behavioural testing was conducted similarly to Cohort 1 in adulthood only (range 74–110), and consisted of OF, NOR, EPM and PPI followed by *in vivo* structural and diffusion MRI (note that MRI was not conducted in adolescence in this cohort). For both cohorts, behavioural testing was conducted 2–7 days before the adulthood scan.

The sample size of Cohort 1 was statistically powered to detect medium effect sizes in regional volume using TBM across four groups (males and females analysed separately), with a minimum sample size of n = 15 per group based on previously observed variance with this method by our group (Serrano et al. 2023). Cohort 2 sample size was powered to detect medium effect sizes in regional TBM with sexes combined, n = 16 per group.

### In vivo MRI

Two to three days after behavioural testing, mice were imaged using a Bruker BioSpec 9.4 T scanner with an 86-mm volume resonator for transmission and a 4-channel surface array coil. Anaesthesia was induced with 4% isoflurane in medical air (1 L/min) and oxygen (0.4 L/min), maintained at 2% but adjusted based on respiration rates. For BOLD fMRI in Cohort 1, we used a medetomidine and isoflurane anaesthesia optimised for mouse fMRI (Grandjean et al. 2014). This consisted of a subcutaneous medetomidine bolus (0.05 mg/kg) followed ten minutes later by its continuous infusion (0.1 mg/kg/h), with isoflurane levels gradually reduced to 0.45-0.65% over 15 minutes from the start of the infusion. BOLD fMRI was conducted after the structural scans, which took a further 45-60 minutes after reducing isoflurane level. The respiration rate was monitored with a pressure sensor, and temperature was monitored with a rectal thermometer and maintained at 36– 37°C using a water circulation system.

### Ex vivo MRI

Following the adulthood *in vivo* scan in Cohort 1, mice were perfused transcardially with 20 mL phosphate-buffered saline (PBS) followed by 4% paraformaldehyde (PFA). Heads were stored in PFA for 48 hours, then transferred to PBS containing 0.05% sodium azide and 2 mM gadolinium-based contrast agent (Gd-DO3A-butrol). Brains were scanned *in cranio* in groups of four using a custom-made holder immersed in perfluoropolyether (Galden®, Solvay).

### MRI acquisition parameters

For *in vivo* imaging, we first acquired an Actual Flip Angle Imaging (AFI) sequence for B1 mapping. Then we acquired three types of 3D multi-gradient-echo images that were used for creating study-specific templates: magnetization-transfer weighted (MTw), proton-density weighted (PDw), and T1-weighted (T1w). Subsequently, T2-weighted Rapid Acquisition with Relaxation Enhancement (RARE) images were obtained. Diffusion-weighted images were then acquired using a single-shot spin-echo echo planar imaging (EPI) sequence. Finally, for the longitudinal MRI study, BOLD rsfMRI data were acquired using a single-shot gradient-echo EPI sequence with 720 repetitions. Additionally, spin-echo EPI image pairs with opposing phase-encoding polarity were recorded to enable correction of susceptibility-induced distortions. The *in vivo* scanning session lasted 1-1.5 hours.

For *ex vivo* morphometric analysis, 3D T2-weighted images were acquired using RARE sequences, with a total scan duration of 1 hour and 5 minutes. *Ex vivo* diffusion-weighted images were obtained using Stejskal-Tanner pulsed gradient spin-echo sequences with a 3D segmented EPI readout. Three b0 images were collected at the beginning of three blocks of 30 diffusion-weighted images. The total scan time for this acquisition was 14 hours and 15 minutes.

### Structural MR image processing

For preprocessing MTw, T1w, and PDw images, de-ringing was conducted using the MRtrix3’s (Tournier et al. 2019) mrdegibbs command. MTw, T1w, and PDw images were averaged across echo times, rigidly co-registered using Advanced Normalization Tools (Avants et al. 2011) antsRegistration, and used for template construction (see below).

For DWI, MRtrix3 dwidenoise was used for de-noising, MRtrix3 mrdegibbs for de-ringing, and FSL’s (Jenkinson et al. 2012; Smith et al. 2004) topup and eddy for susceptibility and eddy-current distortion and motion correction. FSL’s dtifit was used for diffusion tensor imaging (DTI) model fitting, enabling the calculation of fractional anisotropy (FA), mean diffusivity (MD) and axial diffusivity (AD), the latter two which are not reported in this manuscript for brevity. For a more detailed diffusion processing protocol see (Kim et al. 2023).

### Study templates

The antsMultivariateTemplateConstruction2.sh script from ANTs was used to create study specific templates from processed images. For *in vivo* scans of Cohort 1, MTw, T1w, R2* map (generated from the multi-gradient-echo PDw, T1w and MTw images using the qi mpm_r2s command in the QUIT package), S0, FA and MD images were used. For the *ex vivo* scans of Cohort 1, separate T2w and DTI (comprising S0, FA, and MD images) templates were created. For Cohort 2, PDw, T1w, MTw, S0 (estimated non-diffusion-weighted image from dtifit), FA, and MD images were used.

### Jacobian determinant maps

To estimate volume, Jacobian determinant maps were generated from the deformation fields corresponding to the transformation of each subject to the study template using the CreateJacobianDeterminantImage command from ANTs. The Jacobian determinant values of all voxels within the template brain mask were summed to obtain the total brain volume of each subject. Jacobian determinants were calculated from the combined rigid, affine, and Symmetric Normalization (SyN) transforms as well as from only the SyN transforms to obtain maps of absolute and relative volume (accounting for differences in global brain volume), respectively. For TBM, the Jacobian determinants were subsequently log-transformed.

### Voxel-wise analysis

For voxel-wise statistics, FSL randomise with permutation testing was used (10,000 for Cohort 1, 5000 iterations for Cohort 2) followed by a threshold-free cluster enhancement (TFCE) and family-wise error (FWE) correction as described in (Wood et al. 2016; Kim et al. 2023). Given the absence of genotype differences in total brain volume, TBM regional volumes are reported relative to total brain size for greater accuracy (Lerch et al. 2012), while absolute volume maps are reported in the Supplementary files where sex differences in total brain volume were present.

### Common coordinate space

The study template was registered to the Allen Mouse Brain Common Coordinate Framework (Wang et al. 2020) using ANTs, and the Allen atlas was subsequently transformed to the study template space with the inverse transform.

### ROI-based analysis of volume

The images were segmented using an in house modified version of the Allen atlas of 72 regions (Serrano et al. 2023; Wang et al. 2020). These segmentations were subsequently used to compute regional volumes by summing the Jacobian determinants within each parcellation.

Regional volume variability was estimated by calculating a coefficient of variation (CV) for each region with normalised root-mean-square method for each genotype in each experiment, using the calculation: *CV* = σ/μ, where σ is the standard deviation and μ the population mean for each region in the atlas (n = 72 regions). Group differences in CV were calculated with a Kruskal-Wallis test (SciPy.stats, kruskal).

### Fractional anisotropy

For mass-univariate voxel-wise analysis of fractional anisotropy, values from dtifit for Cohort 1 were again analysed with FSL randomise with permutation testing (10,000 permutations) followed by TFCE and FWE-correction. FA is not reported for Cohort 2 in this manuscript for brevity. For ROI-based analysis, voxel FA medians within parcellation were used for all white matter regions.

To calculate the change over time in fractional anisotropy in the longitudinal study, adolescence values for each voxel value or regional median were subtracted from adulthood values. Differences between genotypes were calculated with a mixed ANOVA with between-subjects factor of genotype and within-subjects factor of region. Change from 0 was calculated with a two-sided one-sample *t*-test (SciPy.stats, ttest_1samp), which was corrected with the Benjamini–Hochberg method.

### BOLD fMRI pre-processing

Images were largely pre-processed using the Analysis of Functional NeuroImages (AFNI) toolkit. Slice timing correction was performed using the 3dTshift package, despiking with 3dDespike, and motion correction was applied with 3dvolreg. The motion-corrected time average was then registered to each subject’s own T2w image using ANTs, followed by registration to a template T2w image derived from a separate mouse study conducted at the BRAIN Centre (KCL). The use of this external template was justified by the low resolution of fMRI, which does not benefit from creating a study-specific template.

The images were also distortion corrected using FSL’s topup, with the distortion estimated using an auxiliary pair of spin-echo images with opposite phase encoding polarity and otherwise matching acquisition parameters to the gradient echo EPI used for the BOLD signal. Prior to analysis, corrected images were band-pass filtered at 0.01–0.2 Hz with AFNI’s 3dTproject to remove low-frequency scanner drift noise and high-frequency physiological noise. Nuisance variables (movement and CSF signal) were also simultaneously regressed out of the signal at this stage. Finally, spatial smoothing was applied using AFNI’s 3dBlurInMask with a FWHM kernel.

### Functional connectivity and graph theory analysis

For functional analysis, a high-level parcellation scheme was applied to segment 36 unilateral regions (18 per hemisphere) excluding white matter from the acquired 3D volume. The BOLD signal time-courses were averaged within each ROI, and the mean time-courses were extracted using FSL’s fslmeants tool. Pearson correlation coefficients were calculated for each time-course pair, producing a 36×36 correlation matrix for each subject. These matrices were analysed as functional connectivity (FC) graphs, with edge strength determined by the Fisher z transformed Pearson correlation coefficient. To identify the strongest connections, graphs were thresholded at 5% intervals from 5% to 50%. At each threshold level, referred to as the graph sparsity interval, connections below threshold were set to zero, generating 10 sparsity graphs per subject. FC was calculated as the average non-zero connectivity at each threshold level. Global graph metrics, global efficiency and clustering coefficient, were computed at each sparsity level using Brain Connectivity Toolbox algorithms (Rubinov and Sporns 2010) implemented with Network X (3.4.2) global_efficiency and clustering passing binary thresholded Pearson matrices to prioritise topology in the presence of noise. For global efficiency and clustering coefficient, random curves were calculated by shuffling the thresholded binary matrix positions. The area under the curve (AUC) was computed per subject using trapezoidal numerical integration (numpy.trapz), and the likelihood of observed values was estimated with permutation testing (10,000 permutations).

To estimate changes over time, global graph metric values at adolescence were subtracted from adulthood values at each sparsity interval. Group differences were again tested by calculating the AUC and applying permutation testing. Changes from baseline (zero) were evaluated using two-sided one-sample *t*-tests (difference from 0), corrected for multiple comparisons with the Benjamini-Hochberg (BH) method.

### FDR correction and network-based statistics

Matrices for *C3ar1*-deficient and wild-type mice were compared with Student’s *t*-tests for each pairwise connection, resulting in 36×36 *t*-statistic and *p*-value matrices. To control for multiple comparisons, FDR correction was applied to the upper triangle of the *p*-value matrix using the BH procedure (statsmodels.stats.multitest.fdrcorrection, α = 0.05).

For network-based statistics (NBS), the above-described *t* matrices were thresholded at |*t*| ≥ 2. To identify connected components within the FC graph, adjacency matrices representing significant connections (|*t*| ≥ 2) were converted into graph objects using NetworkX. Regions of interest (ROIs) were treated as nodes, and significant connections as edges. Connected components were identified using a breadth-first search (BFS) algorithm, which explores all neighbouring nodes before moving deeper into the graph. Only components containing more than one ROI were retained for further analysis. To generate a null distribution of maximal component sizes, group labels were randomly shuffled across subjects for each permutation while preserving matrix structure (10,000 permutation). The *p*-value for an observed component was calculated as the proportion of permutations where the maximal component size exceeded that of the observed component.

### *A priori* node strength analysis

We selected 20 (10 left +10 right) anxiety and fear related regions (cingulate cortex, prefrontal cortex, amygdala, pallidum and nucleus accumbens, striatum, hypothalamus, dorsal hippocampus, ventral hippocampus, periaqueductal grey, brain stem) and calculated their average absolute connectivity to all other regions using Fisher z-transformed Pearson correlation coefficients. We then used a mixed ANOVA with between-subjects factor of genotype and within-subjects factor of region followed by pairwise testing with BH FDR *p* value correction.

### Within-network analysis

Two mouse resting state networks were subset from correlation matrices: the default mode network (DMN) and the salience network (Grandjean et al. 2020; Sforazzini et al. 2014). The DMN included bilateral prefrontal cortex, cingulate cortex, and dorsal hippocampus, while the salience network comprised bilateral cingulate cortex, amygdala, and striatum. Additionally, a third anxiety-related network was defined, consisting of the 20 regions described in the node-strength analysis above.

For each network, mean FC was calculated as the average of all pairwise connections between nodes within the network, without applying a threshold. For individual network global efficiency analysis, thresholding was not used due to the small number of nodes. Instead, Pearson matrices normalised to range from 0-1 were passed to bctpy (0.6.1) efficiency_wei which calculated (not binary) efficiency. Student’s *t* tests were used to test significance, and the comparisons were corrected within outcome measure with the BH method.

### Voxel-wise seed-to-brain analysis

We conducted voxel-wise seed-based FC analyses for anxiety-related regions, as well as the colliculus and sensory cortex, which served as control regions not specific to anxiety. For each seed region, the time-course of the BOLD signal was extracted and regressed with the BOLD signal of every voxel in the brain, resulting in a 3D spatial map of the connectivity with the seed. Group-level comparisons of these maps were performed between genotypes using voxel-wise permutation tests with FSL’s randomise (5000 permutation), converted with TFCE and statistical significance corrected for multiple comparisons using FWE (seed-to-brain), but they were not corrected for the presence of multiple seeds.

### General behavioural procedures

For Cohort 1, EPM was administered as the first test, followed by OF. For Cohort 2, OF was the first test, followed by NOR test after two days of low light habituation (4 lux), EPM, and PPI.

Handling of the mice began 2–3 days prior to the behavioural testing battery. By the start of the experiments, the mice sat comfortably on the experimenter’s hand. Mice were handled using cardboard tunnels to minimise stress, and tail handling was avoided.

Behavioural tests were conducted during the dark phase (between 7:00 PM and 11:00 PM) of the light-dark cycle to align with the active period of mice. Mice were randomised by genotype and counter-balanced by sex, with the experimenter systematically blinded to genotype throughout testing and analysis though the allocation of a study ID and test order ID, respectively.

For cleaning of the test apparatus, we used 70% EtOH for all arenas except for the EPM arena for which we used Virusolve (Amity International) to avoid damage to the material. Behaviour was recorded using a Google Pixel 5a camera at 1080p/60 fps. Videos were then cropped and down-sampled using FFmpeg.

### Open field test

Mice were placed in a 40×40×40 cm white arena and allowed to explore freely. For Cohort 1, dim red light (4 lux) was used, while for Cohort 2, the OF test was conducted under bright overhead lighting (500 lux). 10-minute videos were analysed using CleverSys (VA, USA). Outcome parameters included time spent in the centre, total distance travelled, velocity, and thigmotaxis (edge exploration).

### Novel object recognition

Cohort 2 mice were initially habituated to the arena under low light conditions (4 lux) over two days, five minutes per session. On the training day, two identical objects were introduced, and mice were allowed to explore for five minutes. After a one-hour delay, one object was replaced with a novel one. The videos were analysed using CleverSys software stereotypic event “Sniffing” module. Objects were manually outlined with the polygon tool. An interaction with an object was recorded when the mouse’s nose was within 5 mm of the object. The novelty preference was determined by calculating the proportion of time spent exploring the novel object relative to the total exploration time of both objects, with a recognition index chance level of 50%.

### Elevated plus maze

For Cohort 1, EPM was administered to behaviourally naïve animals, while for Cohort 2, it was conducted after OF and NOR tests. In both cases, mice were placed in the closed arm of the arena (65×65×55 cm, elevated 40 cm) under full overhead lighting (500 lux) and allowed to explore for five minutes. The arena was divided into closed, middle, and open areas for analysis with CleverSys software. The number of head dips and stretch-attend postures was recorded using BORIS software, and testing accuracy was compared with the results of an independent scorer.

### Prepulse inhibition

Like NOR, PPI was only conducted in Cohort 2. We used an acoustic startle chamber (SR-LAB, San Diego Instruments, San Diego, CA, USA) with a cylindrical Plexiglas enclosure horizontally mounted on a mobile platform within a sound-proofed isolation chamber. A high-frequency loudspeaker positioned above the enclosure emitted continuous background noise at 65 dB, along with the experimental acoustic stimuli. The startle response was recorded by converting Plexiglas enclosure vibrations into millivolt signals using a piezoelectric unit.

Each session started with a five-minute acclimatisation period to the 65 dB background noise, followed by five startle-alone trials at 120 dB to ensure habituation. Mice then received 10 prepulse stimuli at 3, 6, and 12 dB above background, each preceding a 120 dB pulse in a pseudo-randomised order, interspersed with 10 no-stimulus trials and 10 startle-alone trials. The session concluded with five final startle-alone pulses. The inter-trial interval was randomised between 9-15 seconds to prevent expectation-based modulation of the startle response. %PPI was calculated for each prepulse using the formula: 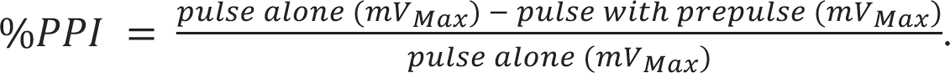

### Statistical procedure for behavioural outcome measures

All statistical analyses were conducted using Python 3.11.7, using relevant libraries such as SciPy and statsmodels. For both studies, the Shapiro-Wilk test was used to assess the normality of data distributions, while Levene’s test was used to evaluate homogeneity of variances. When either assumption of normality or equal variance was violated, the Kruskal-Wallis test was used, followed by Dunn’s test for post hoc pairwise comparisons with Bonferroni correction for multiple testing where applicable. For datasets meeting parametric assumptions, different approaches were used based on the study design. For Cohort 1, a two-way ANOVA was performed to analyse the effect of sex, genotype and sex-by-genotype interaction, followed by Tukey’s test for post hoc comparisons. Bonferroni correction was applied to adjust for multiple comparisons. Due to the relatively small sample size (n = 38) in Cohort 2, sex-by-genotype interactions were not analysed. Instead, Student’s *t*-test was used for comparisons between groups.

To balance statistical rigor with preserving power in exploratory contexts, behavioural measures were not universally corrected for multiplicity. Instead, robustness was inferred through replication across independent cohorts and broad phenotypic consistency. The threshold for statistical significance was set at *p* < 0.05. Individual mice served as the experimental units in all analyses.

## Code availability

Data and code to reproduce figures will be added to https://github.com/hannalemmik upon publication. The MRI processing code is available upon request from Eugene Kim.

## Data availability

Behaviour videos will be added to figshare, and MRI data will be added to openneuro. The intermediary analysis files will be added to github at https://github.com/hannalemmik.

## Acknowledgements

We thank Bao Lu and Craig Gerard for providing the *C3ar1* knockout mice, Marija M. Petrinovic for providing prepulse inhibition testing equipment, KCL Biological Service Unit staff for animal care and François Kroll for comments on this manuscript. This work was funded by the MRC grant “Complement C3aR in adolescent synaptic pruning and risk for anxiety” (MR/W004607/1). Hanna Lemmik was funded by the Wellcome Trust as part of the “Neuro-Immune Interactions in Health & Disease” Wellcome Trust PhD Programme (218452/Z/19/Z).

## Author contributions

HL - writing - original draft, writing - review & editing, conceptualization, methodology, software, formal analysis, data curation, investigation, visualization, funding acquisition; EK - methodology, software, formal analysis, investigation, data curation, writing - review & editing, visualization, funding acquisition; EM - methodology, software, investigation, writing - review & editing; DM - validation, methodology, investigation; MB - software, formal analysis, data curation, visualization; ZL - investigation, validation; DA - investigation, Esther - investigation; MES - funding acquisition; WZ - resources, supervision; AI - resources, supervision; DC - writing - original draft, writing - review & editing, supervision, project administration, funding acquisition, conceptualization, methodology; LW - writing - original draft, writing - review & editing, supervision, funding acquisition, conceptualization, methodology.

