## Supplemental figures and tables for "Complement receptor *C3ar1* deficiency does not alter brain structure or functional connectivity across early life development"

### Supplementary figures and tables

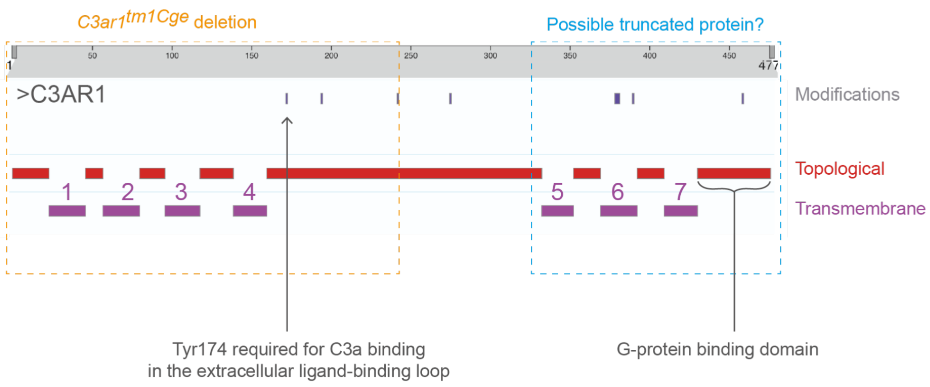


**Supplemental figure 1 | Schematic of the C3AR1 protein.** Marked are the tm1Cge deleted region (246 residues), removing the 4 transmembrane domains and the C3a binding residue at Tyr174. There is an alternative start codon 80 residues downstream of the mutation, meaning that a truncated protein that retains the three last transmembrane domains and the G-protein binding region is possible.

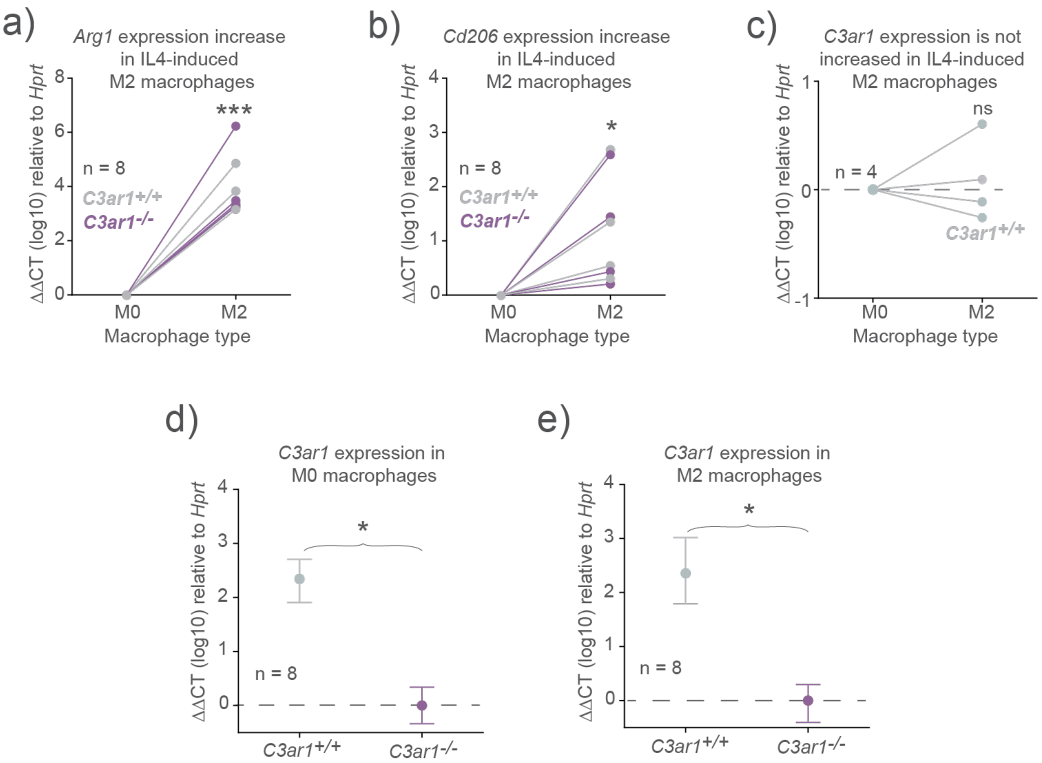


**Supplemental figure 2 | *C3ar1$^{tmCge/tmCge}$* M0 and M2 macrophages do not express *C3ar1*.** qPCR of cDNA extracted from *C3ar1$^{+/+}$* and *C3ar1$^{-/-}$* bone marrow derived macrophages. **(a, b)** Adding recombinant interleukin 4 (IL4) to bone marrow derived M0-like macrophages increased the expression of M2-like macrophage markers *Arg1* and *Cd206* (one-sample *t* tests, M2 expression level difference from 0 (M0, baseline), *Arg1* $t_{[7]} = 10.29$, $p < 0.001$, *Cd206* $t_{[7]} = 3.37$, $p < 0.05$. **(c)** No difference in *C3ar1* expression between M0 and M2 macrophages (one-sample *t* test, M2 expression level difference from 0 (M0, baseline), $p$ = ns). **(d, e)** No *C3ar1* RNA in either M0 or M2 bone marrow derived macrophages from *C3ar1$^{-/-}$* mice (Kruskal Wallis test *C3ar1$^{+/+}$* vs *C3ar1$^{-/-}$*, M0 $p < 0.05$, M2 $p < 0.05$). Data are presented as mean ± 95% CI.

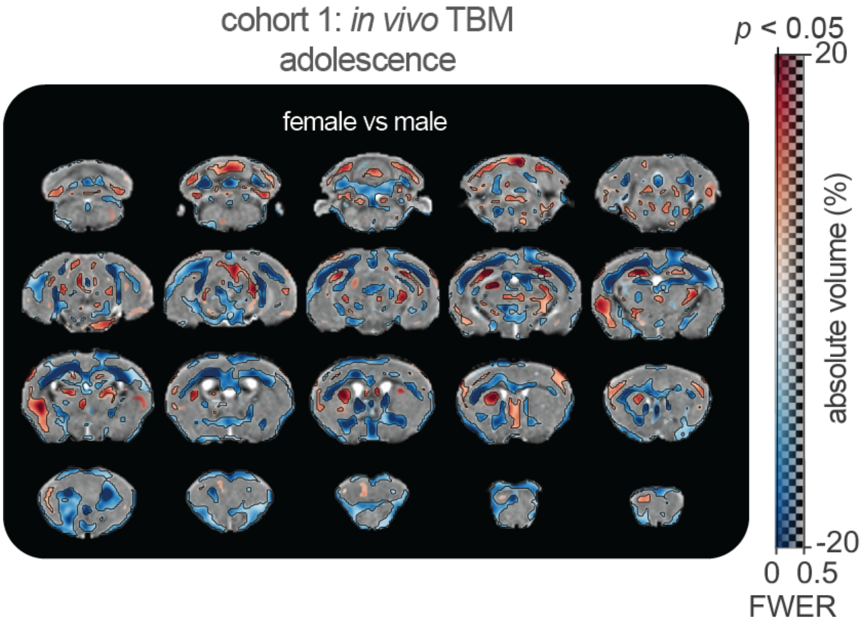


**Supplemental figure 3 | Sex influences regional brain volumes in adulthood:** Cohort 1 *in vivo* absolute volume data. Panel showing absolute regional volume changes (%) overlaid on study-specific coronal template (grey). Red hues signify areas larger in females and blue hues signify areas larger in males . Transparency of the colour overlay shows the statistical significance, ranging from FWE-corrected $p$ value 0.5 to 0 (transparent to opaque, respectively). Areas where FWE-corrected $p$ value < 0.05 are demarcated with a black line, and where the $p$ value > 0.5 are grey (no overlay). Genotypes are combined, males n = 35, 17 *C3ar1*$^{+/+}$ and 18 *C3ar1*$^{-/-}$ ; females n = 34, 18 *C3ar1*$^{+/+}$ and 16 *C3ar1*$^{-/-}$.

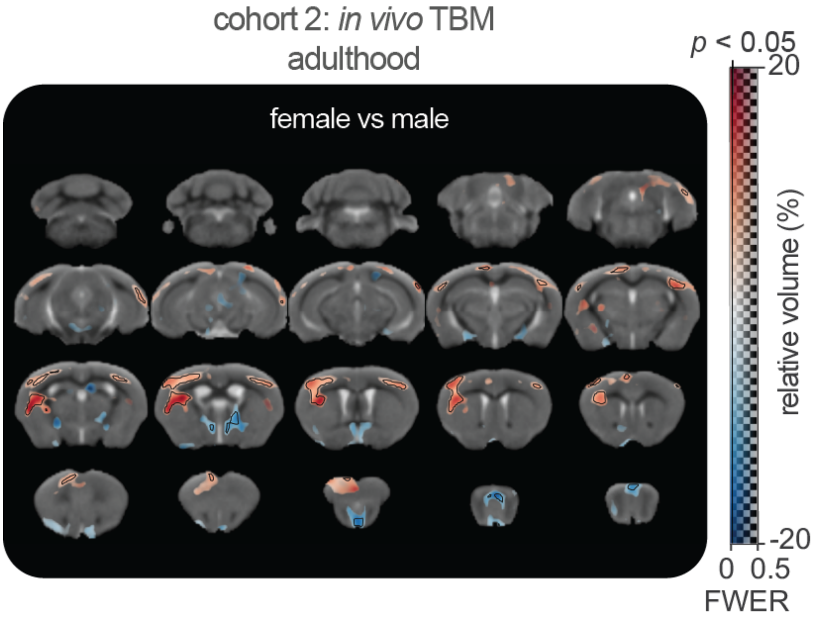


**Supplemental figure 4 | Sex influences regional brain volumes in adulthood:** Cohort 2 *in vivo* data**.** Panels showing relative brain volume changes (%) derived from TBM analysis of *in vivo*. Genotypes are combined, males n = 21, 11 *C3ar1*$^{+/+}$ and 10 *C3ar1*$^{-/-}$; females n = 16, 8 *C3ar1*$^{+/+}$ and 8 *C3ar1*$^{-/-}$.

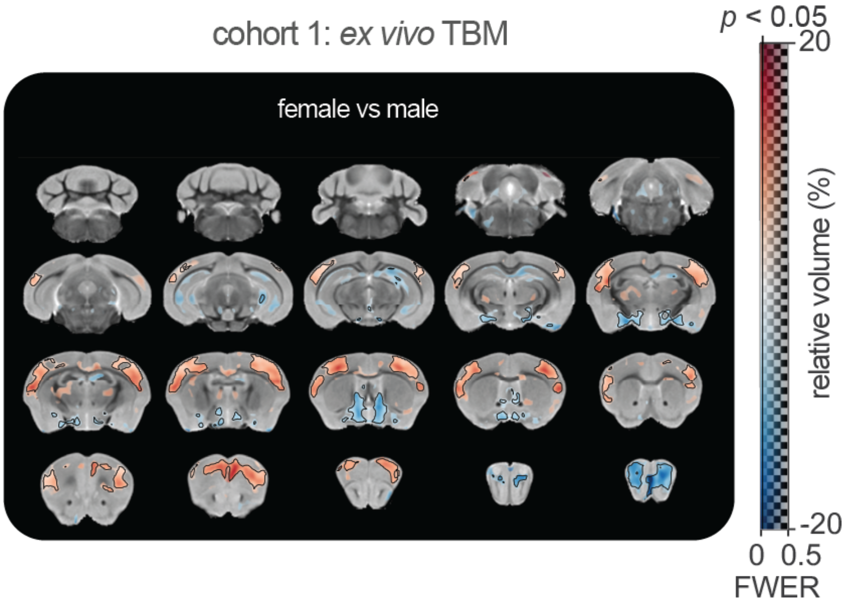


**Supplemental figure 5 | Sex influences regional brain volumes in adulthood**: Cohort 1 *ex vivo* regional volume data. Panels showing TBM relative brain volume changes (%). Genotypes are combined, males n = 35, 17 *C3ar1*$^{+/+}$ and 18 *C3ar1*$^{-/-}$; females n = 34, 18 *C3ar1*$^{+/+}$ and 16 *C3ar1*$^{-/-}$.

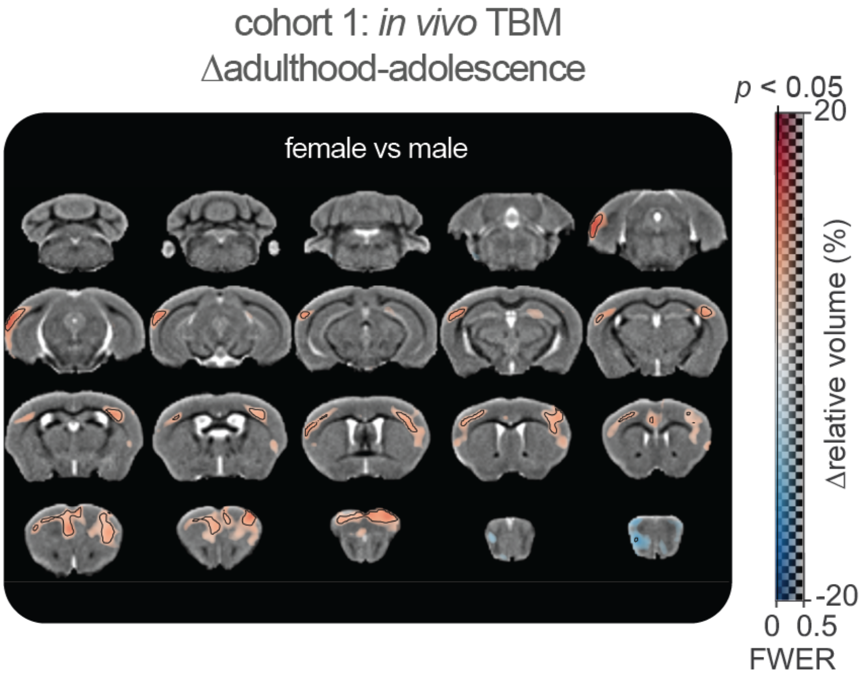


**Supplemental figure 6 | Cortical area relative volume increased more in females than males between PND30 and PND90.** Panels showing change in relative brain volume (Δadulthood – adolescence) between sexes. Genotypes are combined, males n = 35, 17 *C3ar1*$^{+/+}$ and 18 *C3ar1*$^{-/-}$; females n = 34, 18 *C3ar1*$^{+/+}$ and 16 *C3ar1*$^{-/-}$.

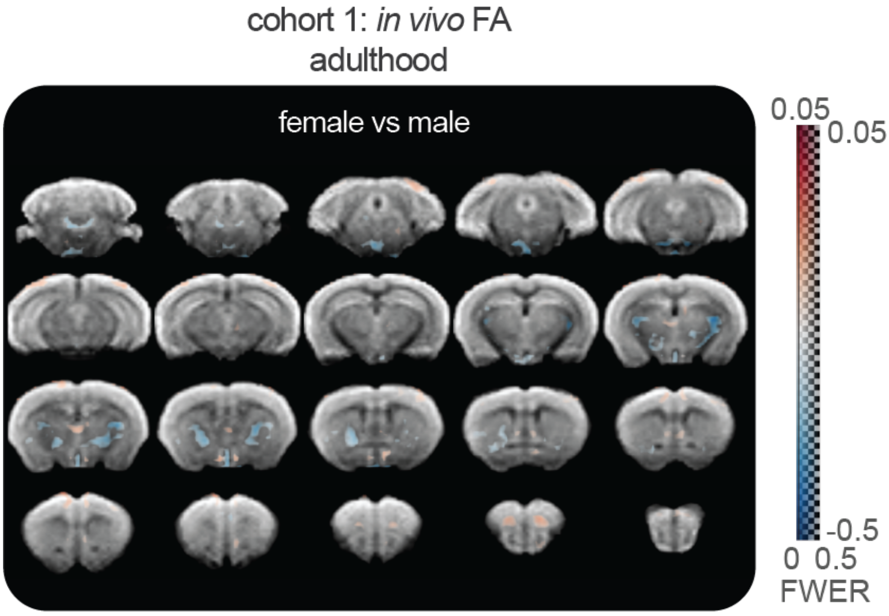


**Supplemental figure 7 | Females show subthreshold decreases in FA at PND90 in voxel-wise fractional anisotropy analysis** corrected for FWE. There are no significant voxels (no black contour) where sex effect is significant ($p < 0.05$). Genotypes are combined, males n = 35, 17 *C3ar1*$^{+/+}$ and 18 *C3ar1*$^{-/-}$; females n = 34, 18 *C3ar1*$^{+/+}$ and 16 *C3ar1*$^{-/-}$.

**Supplemental table 1 | Global functional connectivity metric *p* values with sexes combined.** *C3ar1*$^{-/-}$ vs *C3ar1*$^{+/+}$ permutation testing (10,000 iterations) uncorrected *p* values and Benjamini-Hochberg connected *p*-values (n = 3 *p*-values per metric). FC = functional connectivity, CC = clustering coefficient, GE = global efficiency, PND = postnatal day, BH = Benjamini-Hochberg.

| Metric | *p*-value | BH *p*-value |
|---|---|---|
| FC PND30 | 0.1045 | 0.3135 |
| FC PND90 | 0.3587 | 0.53805 |
| FC change | 0.6282 | 0.6746 |
| GE PND30 | 0.0214 | 0.1926 |
| GE PND90 | 0.2092 | 0.4707 |
| GE change | 0.6746 | 0.6746 |
| CC PND30 | 0.0925 | 0.3135 |
| CC PND90 | 0.3535 | 0.53805 |
| CC change | 0.6638 | 0.6746 |

**Supplemental table 2 | Global functional connectivity metric *p* values with sexes separated.** Permutation testing (10,000 iterations) uncorrected *p* values. AUC = area under the curve (sparsities), KO = *C3ar1*$^{-/-}$, WT = *C3ar1*$^{+/+}$, M = male, F = female, FC = functional connectivity, CC = clustering coefficient, GE = global efficiency, PND = postnatal day.

| Metric | Comparison | Difference in AUC | *p*-value |
|---|---|---|---|
| FC PND30 | M KO vs M WT | 3.6 | 0.15 |
| | M KO vs F KO | 2.05 | 0.44 |
| | M WT vs F WT | 0.49 | 0.85 |
| | F KO vs F WT | 2.05 | 0.44 |
| FC PND90 | M KO vs M WT | 2 | 0.48 |
| | M KO vs F KO | 2.97 | 0.29 |
| | M WT vs F WT | 2.33 | 0.41 |
| | F KO vs F WT | 1.36 | 0.63 |
| CC PND30 | M KO vs M WT | 1.98 | 0.18 |
| | M KO vs F KO | 0.93 | 0.55 |
| | M WT vs F WT | 0.39 | 0.79 |
| | F KO vs F WT | 1.44 | 0.35 |
| CC PND90 | M KO vs M WT | 1.08 | 0.52 |
| | M KO vs F KO | 1.61 | 0.35 |
| | M WT vs F WT | 1.50 | 0.37 |
| | F KO vs F WT | 0.98 | 0.57 |
| GE PND30 | M KO vs M WT | 1.72 | 0.06 |
| | M KO vs F KO | 0.51 | 0.58 |
| | M WT vs F WT | -0.06 | 0.95 |
| | F KO vs F WT | 1.15 | 0.21 |
| GE PND90 | M KO vs M WT | 1.25 | 0.28 |
| | M KO vs F KO | 1.64 | 0.16 |
| | M WT vs F WT | 1.08 | 0.35 |
| | F KO vs F WT | 0.69 | 0.55 |

**Supplemental table 3 | Mean absolute anxiety-related region seed connectivity ANOVA *p* values with sexes separated.** DF1 = degrees of freedom (numerator), DF2 = degrees of freedom (denominator), MS = mean squares (average variance explained), F = F statistic, *p*-unc = uncorrected *p* value, np2 = $\eta_p^2$ (partial eta squared effect size), PND = postnatal day.

| Age | Effect name | DF1 | DF2 | MS | F | *p*-unc | np2 |
|---|---|---|---|---|---|---|---|
| PND90 | group | 3 | 61 | 0.33 | 1.0 | 3.97E-01 | 0.05 |
| | ROI | 19 | 1159 | 0.78 | 174.65 | 1.48e-323 | 0.74 |
| | Group-by-ROI | 57 | 1159 | 0.01 | 1.33 | 5.37E-02 | 0.06 |
| PND30 | group | 3 | 60 | 0.37 | 1.5 | 0.22 | 0.07 |
| | ROI | 19 | 1140 | 0.59 | 138.68 | 4.50E-280 | 0.7 |
| | Group by-ROI | 57 | 1140 | 0.01 | 0.66 | 0.98 | 0.03 |

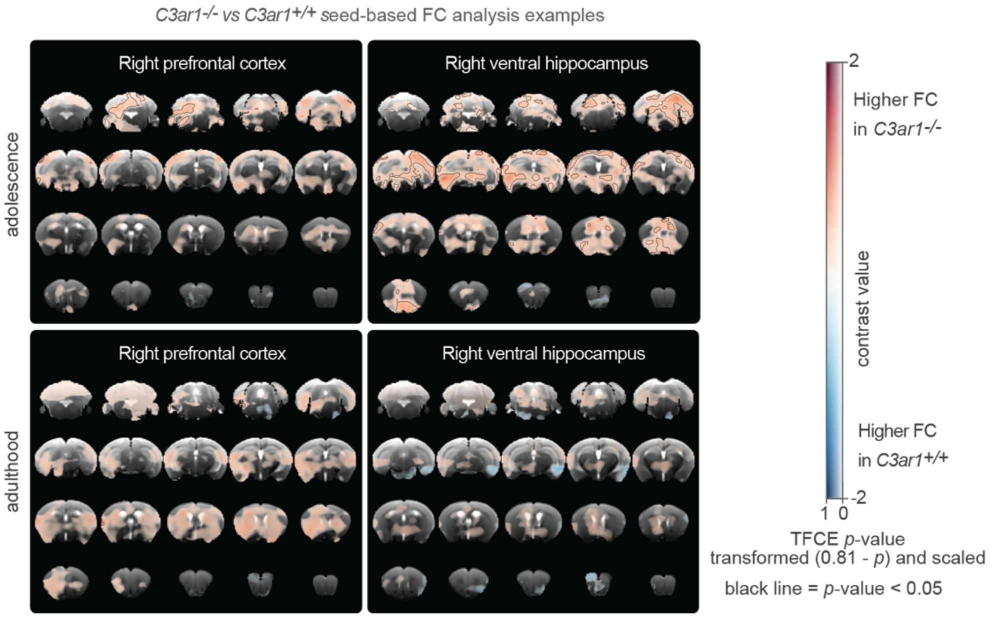


**Supplemental figure 8 | Seed analysis examples in the right hemisphere.** Examples of seed-based FC maps showing voxel-wise group differences between *C3ar1*$^{-/-}$ and *C3ar1*$^{+/+}$ mice (using *t* tests) with seeds placed in the right ventral hippocampus and right prefrontal cortex in adolescence and adulthood datasets. The dual scale bar displays contrast value on the x-axis and threshold free cluster enhancement (TFCE) *p*-values (transformed 0.81-p) on the y-axis. The transformed *p*-values have been re-scaled to range from 0 to 1 for visualisation, with darker colours representing greater statistical significance. Black outlines demarcate regions where TFCE *p*-values are below 0.05.

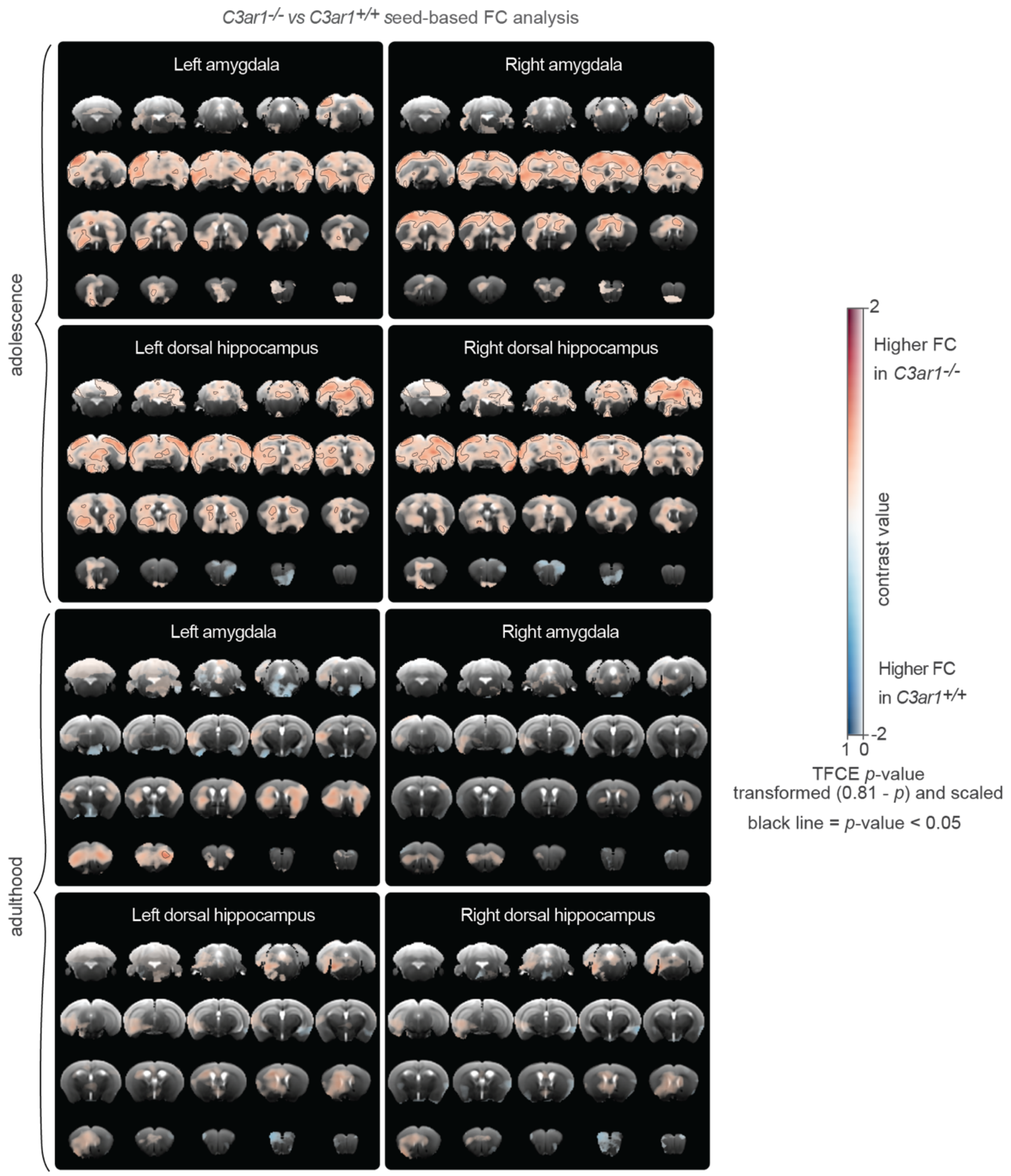


**Supplemental figure 9 | Seed analysis of anxiety-related regions.** Examples of seed-based FC maps showing voxel-wise group differences between *C3ar1*$^{-/-}$ and *C3ar1*$^{+/+}$ mice (using *t* tests) with seeds placed in bilateral dorsal hippocampus and amygdala in adolescence and adulthood datasets. The dual scale bar displays contrast value on the x-axis and threshold free cluster enhancement (TFCE) *p*-values (transformed 0.81-p) on the y-axis. The transformed *p*-values have been re-scaled to range from 0 to 1 for visualisation, with darker colours representing greater statistical significance. Black outlines demarcate regions where TFCE *p*-values are below 0.05.

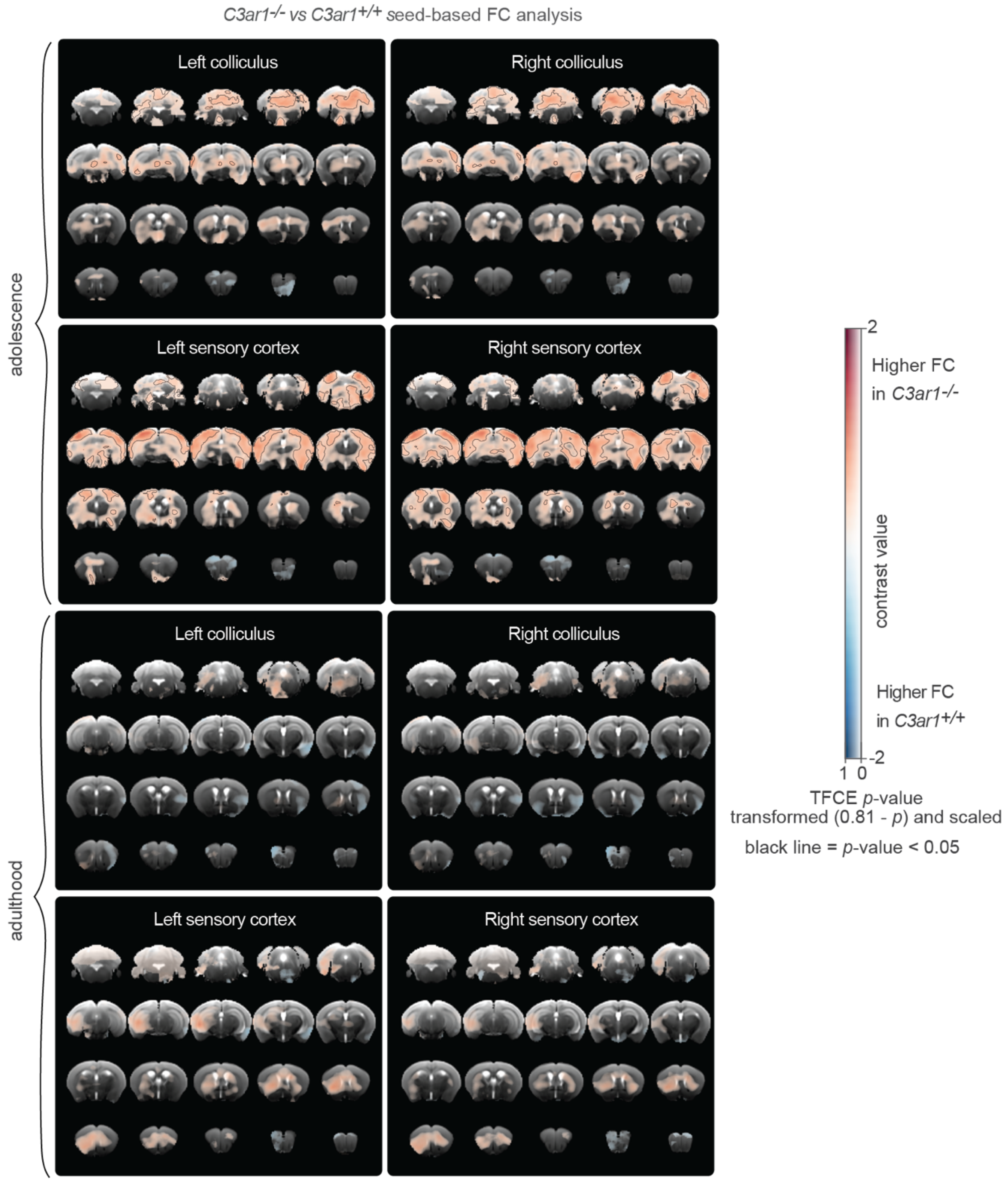


**Supplemental figure 10 | Seed analysis of colliculus and sensory cortex connectivity.** Seed-based FC maps showing voxel-wise group differences between *C3ar1*$^{-/-}$ and *C3ar1*$^{+/+}$ mice (using *t* tests) with seeds placed in bilateral colliculus and sensory in adolescence and adulthood datasets. The dual scale bar displays contrast value on the x-axis and threshold free cluster enhancement (TFCE) *p*-values (transformed 0.81-p) on the y-axis. The transformed *p*-values have been re-scaled to range from 0 to 1 for visualisation, with darker colours representing greater statistical significance. Black outlines demarcate regions where TFCE *p*-values are below 0.05.

**Supplemental table 4 | Anxiety-like behavioural outcome measures**

| Outcome measure | Comment |
|---|---|
| Down with anxiety | |
| OF duration core (30% of arena) | Commonly used metric to quantify anxiety-like behaviour. Similar measurements in *C3ar1*$^{-/-}$ mice were previously reported by (Pozo-Rodrigálvarez et al. 2021) (25% of arena), (Westacott et al. 2022) (36% of arena) and (Sun et al. 2024) (50% of arena), with none finding significant differences compared to control.<br>Open field centre duration is not always sensitive to anxiolytic or anxiogenic drugs (Prut and Belzung 2003). |
| OF duration centre (70% of the arena) | Like the above but allows for more data to be gathered where exploration of the core 30% is insufficient. |
| EPM duration open | Increased with anxiolytics in rats (Pellow et al. 1985) where it is regulated by the ventral hippocampus and the lateral septum (Trent and Menard 2010). Tested in *C3ar1*$^{-/-}$ mice by Westacott et al. (2022) who saw decreased duration relative to wild-type, and Sun et al. (2024) who did not. |
| EPM duration middle | Rodents naturally seek enclosed spaces for protection. In the EPM tests, they first tend to remain in closed arms, then may pause in the middle section to assess safety before entering open areas. Higher anxiety would presumably be associated with more time spent also in this middle zone. |
| EPM bouts open | Like EPM duration open, also specifically validated in rats (Pellow et al. 1985). |
| EPM head dips | Anxiolytic drugs decrease head dip number and duration in the hole-board test in mice (Takeda, Tsuji, and Matsumiya 1998). Tested in *C3ar1*$^{-/-}$ mice by (Westacott et al. 2022) who saw decreased duration relative to wild-type. |
| Up with anxiety | |
| OF duration periphery | Thigmotaxis is a measure of anxiety-like behaviour in mice (Simon, Dupuis, and Costentin 1994). |
| OF latency to core 30% | Similar to core duration, mice who take longer to explore the centre can be more fearful. |

| | |
|---|---|
| OF latency to centre 70% | Similar to the above. |
| EPM latency to open arm | Tested in *C3ar1*$^{-/-}$ mice by Westacott et al. (2022) who saw longer latency relative to wild-type. |
| EPM stretch attend postures | Westacott et al. (2022) saw an increase in *C3ar1*$^{-/-}$ mice. |
| EPM duration closed | Like thigmotaxis. Westacott et al. (2022) saw an increase in *C3ar1*$^{-/-}$ mice. |

**Supplemental table 5 | Uncorrected *p* values for Cohort 1 behavioural measures.**

| Outcome measure | Test type | Geno-type *F* stat | Geno-type *p* value | Sex *F* stat | Sex *p* value | Inter-action *F* stat | Inter-action *p* value | *H* stat | 4 groups *p* value |
|---|---|---|---|---|---|---|---|---|---|
| EPM bouts open | Kruskal | | | | | | | 2.75 | 0.43 |
| EPM head dips | Kruskal | | | | | | | 1.37 | 0.71 |
| EPM duration closed | ANOVA | 0.38 | 0.54 | 0.16 | 0.69 | 2.85 | 0.10 | | |
| EPM duration middle | ANOVA | 0.03 | 0.87 | 1.60 | 0.21 | 0.92 | 0.34 | | |
| EPM duration open | Kruskal | | | | | | | 0.37 | 0.95 |
| EPM latency middle | Kruskal | | | | | | | 0.93 | 0.82 |
| EPM latency open | Kruskal | | | | | | | 1.94 | 0.59 |
| EPM stretch attend postures | ANOVA | 0.68 | 0.41 | 3.90 | 0.05 | 0.94 | 0.34 | | |
| OF distance centre | Kruskal | | | | | | | 0.76 | 0.86 |
| OF distance core | Kruskal | | | | | | | 0.20 | 0.98 |

| | | | | | | | | | |
|---|---|---|---|---|---|---|---|---|---|
| OF distance periphery | ANOVA | 0.49 | 0.49 | 2.09 | 0.15 | 0.00 | 0.98 | | |
| OF duration centre | ANOVA | 1.28 | 0.26 | 0.16 | 0.69 | 0.50 | 0.48 | | |
| OF duration core | Kruskal | | | | | | | 0.55 | 0.91 |
| OF duration periphery | ANOVA | 1.28 | 0.26 | 0.17 | 0.69 | 0.50 | 0.48 | | |
| OF latency centre | Kruskal | | | | | | | 0.33 | 0.95 |
| OF latency core | Kruskal | | | | | | | 1.97 | 0.58 |
| OF loco speed | ANOVA | 0.08 | 0.77 | 5.53 | 0.02 | 0.04 | 0.85 | | |
| OF loco total distance | ANOVA | 0.02 | 0.89 | 3.30 | 0.07 | 0.00 | 0.96 | | |
| OF loco total duration | ANOVA | 0.26 | 0.61 | 1.74 | 0.19 | 0.00 | 0.96 | | |

**Supplemental table 6 | Uncorrected *p* values and effect sizes for Cohort 2 behavioural measures.**

| Outcome measure | n WT | n KO | Para-metric? | Test type | Test stat | *p* value | Effect size type | Effect size |
|---|---|---|---|---|---|---|---|---|
| ASR avg startle | 19 | 20 | FALSE | Kruskal | 0.71 | 0.399 | $\varepsilon^2$ | -0.01 |
| ASR baseline startle | 19 | 20 | FALSE | Kruskal | 0.03 | 0.866 | $\varepsilon^2$ | -0.03 |
| ASR habituation | 19 | 20 | TRUE | *t* test | -1.95 | 0.059 | Cohen's *d* | -0.63 |
| ASR latency | 19 | 20 | FALSE | Kruskal | 0.97 | 0.325 | $\varepsilon^2$ | -0.001 |
| ASR PPI global | 19 | 20 | TRUE | *t* test | 1.42 | 0.163 | Cohen's *d* | 0.46 |
| EPM bouts open | 18 | 18 | FALSE | Kruskal | 0.41 | 0.523 | $\varepsilon^2$ | -0.02 |
| EPM duration closed | 18 | 18 | TRUE | *t* test | 0.44 | 0.662 | Cohen's *d* | 0.15 |
| EPM duration middle | 18 | 18 | TRUE | *t* test | 0.41 | 0.688 | Cohen's *d* | 0.14 |
| EPM duration open | 18 | 18 | FALSE | Kruskal | 0.84 | 0.359 | $\varepsilon^2$ | -0.001 |
| EPM head dips | 18 | 18 | FALSE | Kruskal | 0.01 | 0.987 | $\varepsilon^2$ | -0.03 |
| EPM latency middle | 18 | 18 | FALSE | Kruskal | 0.65 | 0.42 | $\varepsilon^2$ | -0.01 |
| EPM latency open | 18 | 18 | FALSE | Kruskal | 0.9 | 0.342 | $\varepsilon^2$ | -0.003 |
| EPM stretch attend postures | 18 | 18 | TRUE | *t* test | 1.08 | 0.288 | Cohen's *d* | 0.36 |

| | | | | | | | | |
|---|---|---|---|---|---|---|---|---|
| NOR test bouts familiar | 19 | 20 | FALSE | Kruskal | 0.56 | 0.472 | $\varepsilon^2$ | -0.013 |
| NOR test bouts novel | 19 | 20 | FALSE | Kruskal | 0.17 | 0.683 | $\varepsilon^2$ | -0.023 |
| NOR test familiar exploration | 19 | 20 | FALSE | Kruskal | 0.11 | 0.736 | $\varepsilon^2$ | -0.024 |
| NOR test latency novel | 19 | 20 | FALSE | Kruskal | 1.74 | 0.187 | $\varepsilon^2$ | 0.02 |
| NOR test loco duration | 19 | 20 | TRUE | *t* test | 1.14 | 0.261 | Cohen's *d* | 0.36 |
| NOR test novel exploration | 19 | 20 | TRUE | *t* test | -0.6 | 0.523 | Cohen's *d* | -0.21 |
| NOR test RI | 19 | 20 | TRUE | *t* test | -0.42 | 0.673 | Cohen's *d* | -0.14 |
| NOR test total exploration | 19 | 20 | TRUE | *t* test | -0.55 | 0.584 | Cohen's *d* | -0.18 |
| NOR train bouts | 19 | 20 | TRUE | *t* test | 0.5 | 0.618 | Cohen's *d* | 0.16 |
| NOR train exploration duration | 19 | 20 | TRUE | *t* test | -0.20 | 0.840 | Cohen's *d* | -0.07 |
| NOR train locomotion duration | 19 | 20 | FALSE | Kruskal | 2.94 | 0.087 | $\varepsilon^2$ | 0.052 |
| OF distance centre | 19 | 20 | TRUE | *t* test | 2.88 | 0.007 | Cohen's *d* | 0.92 |
| OF distance core | 19 | 20 | FALSE | *t* test | 2.30 | 0.129 | $\varepsilon^2$ | 0.04 |
| OF distance periphery | 18 | 20 | TRUE | *t* test | 1.78 | 0.084 | Cohen's *d* | 0.57 |
| OF duration centre | 19 | 20 | FALSE | Kruskal | 0.97 | 0.325 | $\varepsilon^2$ | -0.001 |
| OF duration core | 19 | 20 | FALSE | Kruskal | 1.78 | 0.182 | $\varepsilon^2$ | 0.02 |
| OF duration periphery | 18 | 20 | FALSE | Kruskal | 0.62 | 0.43 | $\varepsilon^2$ | -0.01 |
| OF latency centre | 19 | 20 | FALSE | Kruskal | 0.001 | 0.978 | $\varepsilon^2$ | -0.03 |
| OF latency core | 19 | 20 | FALSE | Kruskal | 0.44 | 0.509 | $\varepsilon^2$ | -0.06 |
| OF loco speed | 19 | 20 | TRUE | *t* test | 1.33 | 0.190 | Cohen's *d* | 0.43 |
| OF loco duration | 19 | 20 | TRUE | *t* test | 1.78 | 0.083 | Cohen's *d* | 0.57 |
| OF total distance | 19 | 20 | TRUE | *t* test | 2.01 | 0.052 | Cohen's *d* | 0.64 |
| PPI 12dB | 19 | 20 | TRUE | *t* test | 0.28 | 0.783 | Cohen's *d* | 0.09 |
| PPI 3dB | 19 | 20 | TRUE | *t* test | 1.58 | 0.122 | Cohen's *d* | 0.51 |
| PPI 6dB | 19 | 20 | TRUE | *t* test | 1.99 | 0.054 | Cohen's *d* | 0.64 |

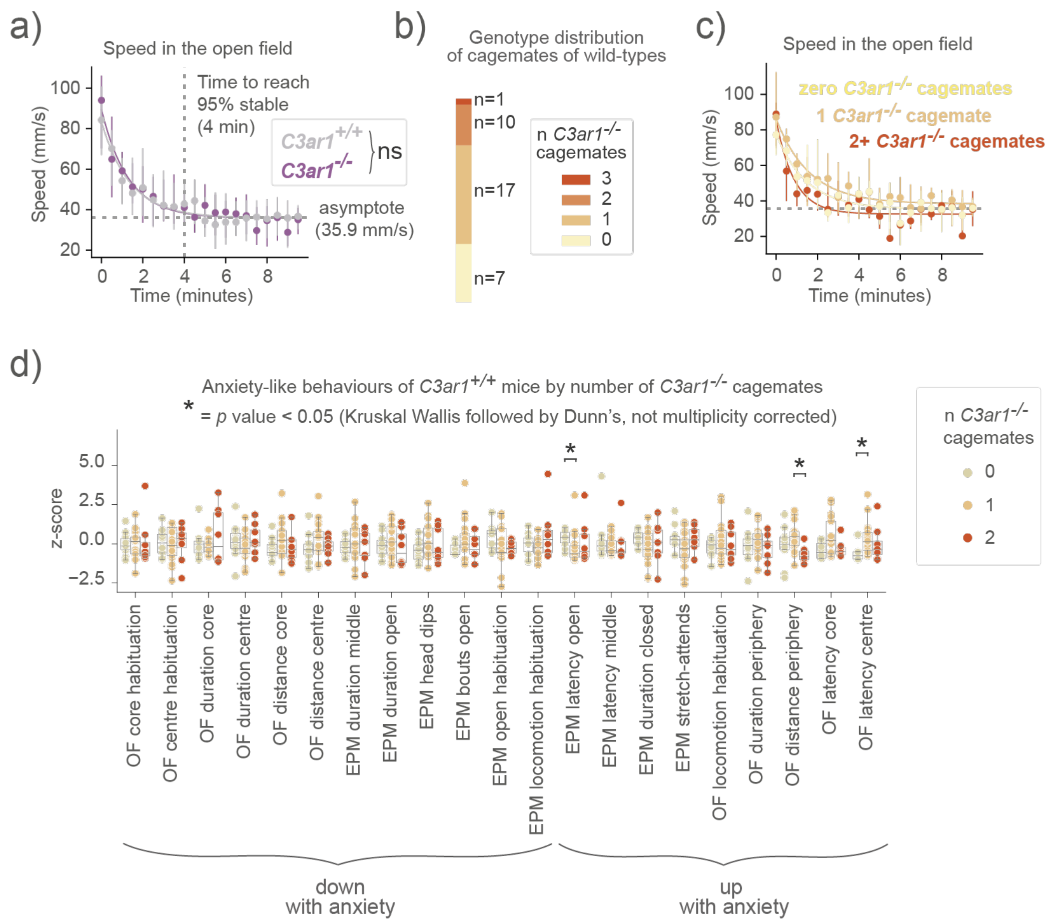


**Supplemental figure 11 | Wild-type anxiety-like behaviour does not depend on the number of co-housed *C3ar1*-deficient mice. (a)** Cohort 1 OF speed in 30-second bins plotted as median and interquartile range with an exponential decay curve. Both groups habituate by the 4th minute (95% asymptote). Two-way ANOVA (time bin, genotype and time-by-genotype interaction, only the effect of time is significant). Shown also are the asymptote and the time animals reach 95% stability in speed **(b)** 35 wild-type mice were housed with either zero, one, two or three *C3ar1*$^{-/-}$ mice. **(c)** Wild-type speed in OF in the longitudinal study cohort separated by the number of *C3ar1*$^{-/-}$ per cage. Two-way ANOVA (time bin, group and time-by-group interaction, only the effect of time is significant). **(d)** Wild-type anxiety-like outcome measures separated by the number of *C3ar1*$^{-/-}$ per cage.
